# A single cell framework identifies functionally and molecularly distinct multipotent progenitors in adult human hematopoiesis

**DOI:** 10.1101/2024.05.07.592983

**Authors:** Asiri Ediriwickrema, Yusuke Nakauchi, Amy C. Fan, Thomas Köhnke, Xiaoyi Hu, Bogdan A. Luca, YeEun Kim, Sreejith Ramakrishnan, Margaret Nakamoto, Daiki Karigane, Miles H. Linde, Armon Azizi, Aaron M. Newman, Andrew J. Gentles, Ravindra Majeti

**Author notes:** Corresponding author and lead contact; address: Lokey Stem Cell Research Building, Room 3005, 265 Campus Dr, Stanford, CA 94305, USA.

## Abstract

Hematopoietic multipotent progenitors (MPPs) regulate blood cell production to appropriately meet the biological demands of the human body. Human MPPs remain ill-defined whereas mouse MPPs have been well characterized with distinct immunophenotypes and lineage potencies. Using multiomic single cell analyses and complementary functional assays, we identified new human MPPs and oligopotent progenitor populations within Lin-CD34+CD38dim/lo adult bone marrow with distinct biomolecular and functional properties. These populations were prospectively isolated based on expression of CD69, CLL1, and CD2 in addition to classical markers like CD90 and CD45RA. We show that within the canonical Lin-CD34+CD38dim/loCD90CD45RA-MPP population, there is a CD69+ MPP with long-term engraftment and multilineage differentiation potential, a CLL1+ myeloid-biased MPP, and a CLL1-CD69-erythroid-biased MPP. We also show that the canonical Lin-CD34+CD38dim/loCD90-CD45RA+ LMPP population can be separated into a CD2+ LMPP with lymphoid and myeloid potential, a CD2-LMPP with high lymphoid potential, and a CLL1+ GMP with minimal lymphoid potential. We used these new HSPC profiles to study human and mouse bone marrow cells and observe limited cell type specific homology between humans and mice and cell type specific changes associated with aging. By identifying and functionally characterizing new adult MPP sub-populations, we provide an updated reference and framework for future studies in human hematopoiesis.

## Introduction

Hematopoiesis, which is the development of differentiated blood cells from a multipotent hematopoietic stem cell (HSC), is a complex, highly regulated, biomolecular process^1^. The careful maintenance of hematopoiesis is critical for meeting the needs of the body through times of stress, infection, and illness. Multipotent progenitors (MPPs) are important regulators of blood cell production and are key for maintaining appropriate cell output over the human lifespan. The definition of human MPPs lacks clarity which in turn limits our understanding of human hematopoiesis. The characterization of blood cells has primarily relied on the prospective isolation of cells using flow cytometry coupled with *in vitro* and *in vivo* functional assays^2,3^. These studies have led to the classical hierarchical tree description of hematopoiesis containing HSCs at the apex capable of long-term self-renewal and multilineage differentiation potential^2,4–7^. Historically, HSCs have been described as long-term HSCs (LT-HSCs) and short-term HSCs (ST-HSCs) based on differential ability for multipotent engraftment upon serial transplantations. The HSC can generate increasingly abundant hematopoietic stem and progenitor cells (HSPCs) and ultimately terminally differentiated blood cells. Historically, HSPC populations have been defined using fluorescence-activated cell sorting (FACS) based on differential cell surface marker expression coupled with *in vitro* and *in vivo* functional assays. The hierarchical tree model has been frequently revised^8^; however, it is believed that HSCs give rise to MPPs, and lymphoid-primed multipotent progenitors (LMPPs), which have decreasing self-renewal capacity but persistent lymphoid and myeloid potential^5,6,9^. MPPs and LMPPs in turn differentiate into increasingly lineage restricted oligopotent and unipotent progenitors. Several oligopotent progenitor (OPP) sub-populations have been described including common myeloid progenitors (CMPs), megakaryocyte-erythroid progenitors (MEPs), granulocyte-monocyte progenitors (GMPs), and common lymphoid progenitors (CLPs or B cell Natural Killer (NK) cell progenitors)^5,6,8,10–12^. The immunophenotypic and functional heterogeneities within OPPs have been re-evaluated leading to a better understanding of the underlying cellular content of these sub-populations^6,13^.

Importantly, human MPPs are functionally and molecularly heterogenous within purified subpopulations using current FACS methods, and they have not been refined based on their immunophenotypic profiles^5,13–15^. This reflects how little is known about the human MPP compartment, especially when compared to the mouse. Studies in murine hematopoiesis have identified distinct MPP subsets^16,17^, which undergo functional changes with stress and aging. Specifically, four murine MPPs, MPP1-4, have been identified: MPP1 is considered a metabolically active HSC subset^18^, MPP2 and MPP3 are considered myeloid-biased MPPs^18^, and MPP4 is considered a lymphoid-biased MPP^19^. Similar immunophenotyping studies in humans are limited and often involve cord blood (CB) cells^5,13,20^, which can be functionally different than adult bone marrow (BM) cells^13,21–24^. Further, studies in both human and mice suggest that early progenitors, defined by traditional immunophenotypes, are lineage primed, and therefore, question the existence of immunophenotypically defined OPPs like the CMP^13,15^.

Although it has been important to use surface marker expression to purify HSPC populations and perform functional assays, this approach has led to the idea that hematopoiesis progresses via a step-wise differentiation pathway^2^. Recent studies reevaluating hematopoiesis using single cell methods challenge this view as they present primitive hematopoiesis as a more gradual and continuous process^7,8,14,25^. These observations are important, but are not radical, as surface marker expression is continuous, and discrete bins of expression are not always present. Regardless, the hierarchical resolution of mouse hematopoiesis using surface markers, particularly the MPP compartment, is finer than in humans and has led to significant advances in our understanding of hematopoiesis. The need remains to resolve primitive human MPPs into distinct, immunophenotypically defined, functionally distinct sub-populations to prospectively isolate and study these cells.

Analysis of single cell RNA-sequencing (scRNA-seq) data from FACS isolated murine HSCs and MPPs suggests that these cells also exist in a continuum despite being immunophenotypically distinct^15,26^. It is therefore important to not only adapt prior methods from the study of mouse hematopoiesis to human HSPCs, but also consider a multi-modal framework to both improve our ability to compartmentalize and study hematopoiesis and associated blood disorders^27,28^. Although multiple scRNA-seq studies in healthy human hematopoiesis have been published over the past 8 years^6,14,29–35^, only a few have incorporated multiple analytes like concurrent scRNAseq and surface marker expression analysis using antibody derived tags (ADTs; scADTseq)^30,34,36^. Importantly, none have functionally examined the heterogeneous populations in the previously defined MPP^5^.

Here, we present a systematic approach for analyzing multiomic single cell data from adult human BM and provide a framework for rigorously evaluating the HSPC compartment of adult hematopoiesis using orthogonal computational methods and functional assays. As a result, we observe greater phenotypic heterogeneity within CD34+ cells in adult BM compared to CB, and newly define 4 functionally distinct progenitors based on the expression of CD90, CD69, and CLL1 within the lineage negative (Lin-) CD34+CD38dim/loCD45RA-CD2-compartment. Using complementary *in vitro* and *in vivo* assays, we observe that 1) the Lin-CD34+CD38dim/loCD90CD45RA-CLL1-CD69+ adult BM population contains a new human MPP with long-term engraftment capacity and 7 lineage differentiation potential, 2) the Lin-CD34+CD38dimCD90CD45RA-CLL1-CD69-population contains an erythroid-biased human MPP and 3) the LinCD34+CD38dimCD90-CD45RA-CLL1+ population contains a myeloid-biased MPP. Both the myeloid and erythroid biased MPPs have significantly decreased engraftment potential *in vivo*. We also observe 3 distinct OPPs within the Lin-CD34+CD38dim/loCD45RA+ adult bone marrow population with differential CD2, CD69, and CLL1 surface marker expression: CD2+ LMPPs with high cytotoxic gene expression and both myeloid and lymphoid potencies, CD2-CLL1-LMPPs with high lymphoid potency, and CD2-CLL+ GMPs with minimal lymphoid potency. Integrated single cell assay for transposase-accessible chromatin using sequencing (scATAC-seq) analysis revealed distinct patterns of chromatin accessibility amongst newly identified HSPCs. Using this updated HSPC profile as a reference, we also performed an integrated cross species analysis which revealed variable similarities between mouse and human HSPCs. Finally, we performed a large-scale (*n*=42,480 cells) computational analysis across 20 normal human donors (ages 0-75) and observe cell type specific transcriptional changes associated with aging and evaluated those changes in common blood cancers. Our work provides additional insights into adult human MPP and OPP biology with implications for future multiomic studies.

## Results

### A multi-modal framework for studying adult human HSPCs

To identify immunophenotypically distinct sub-populations within adult human HSPCs, 10,287 BM mononuclear cells from 3 donors (Figure 1 and Table S1) were profiled using concurrent scRNAseq and scADT-seq (Figure S1, Table S2, Methods). We performed strict quality control to identify high quality cells, and lymphocytes were subsequently removed to specifically study the HSPC and erythro-myeloid compartment (Figure S1A-B; Methods). The subsequent workflow implements 4 key objectives: 1) computational purification of high quality HSPC and myeloid cell clusters, 2) subsequent FACS based purification, functional evaluation, and annotation of newly identified HSPCs, 3) molecular characterization of new HSPCs using multi-dataset integration, 4) cross-species evaluation of human and murine HSPC homology, and 5) large-scale metaanalysis of these new HSPCs in aging and cancer.

**Figure 1:**
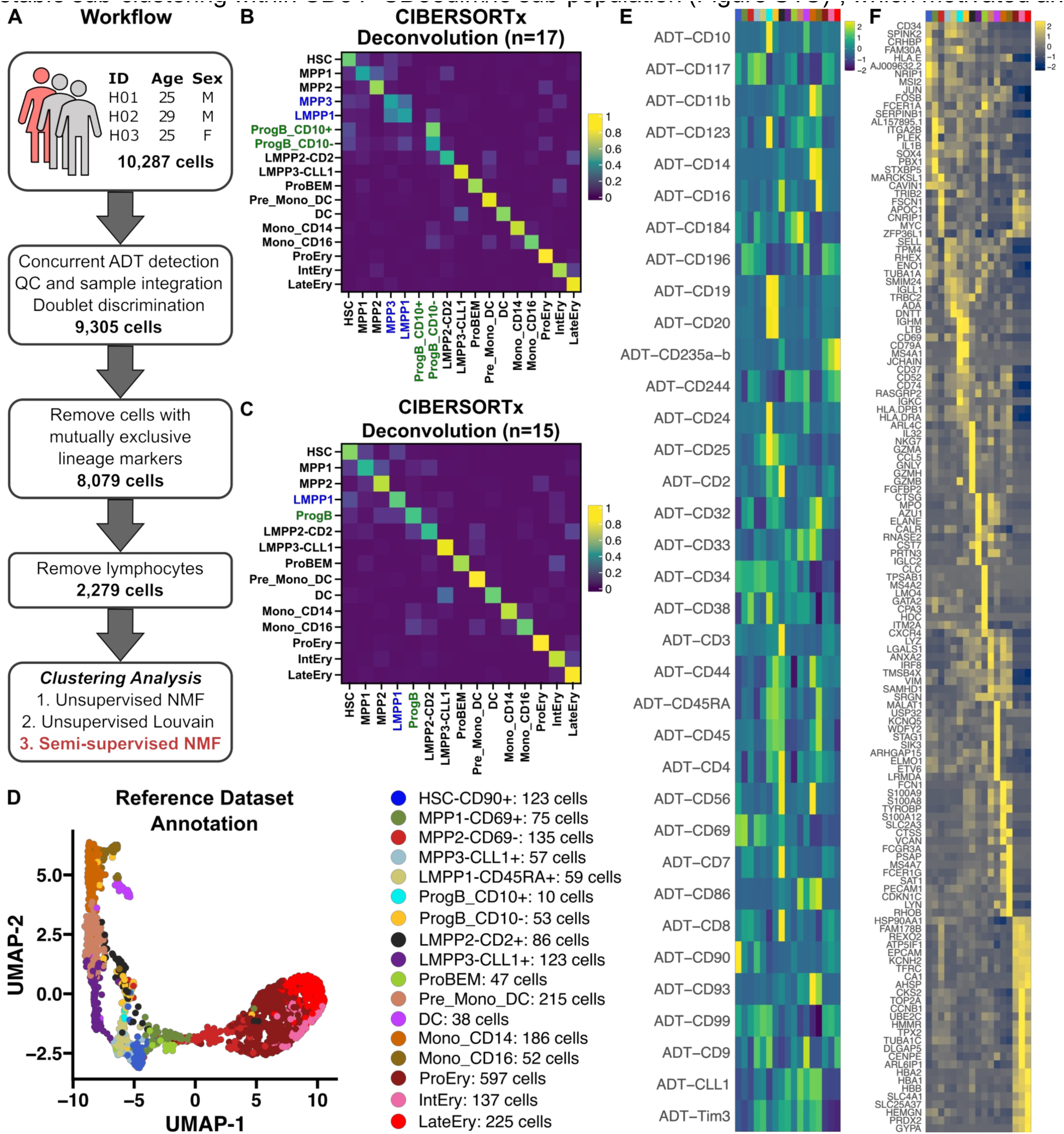
Candidate HSPC sub-populations were identified using NMF. A) schematic illustrating the general workflow for this analysis. B) Heatmap illustrating imputation results from deconvolving artificial bulk transcriptomes derived from cells in each respective cluster. The reference profile was derived from k=18 clusters using NMF with one cluster removed for low cell content. C) Heatmap illustrating imputation results from a similar deconvolution after collapsing transcriptionally similar clusters (see Methods). D) UMAP projection of the reference single cell dataset and final cell counts for each cluster. E) Heatmap showing average ADT expression for each cluster. F) Heatmap showing average gene expression for top features in each cluster. ADT – Antibody derived tags, DC – dendritic cell, EMPP – erythroid-biased MPP, GMP – granulocyte monocyte progenitor, HSC – hematopoietic stem cell, HSPC – hematopoietic stem and progenitor cell, LMPP – lymphoid-primed MPP, MPP – multipotent progenitor, MPPP – myeloid-biased MPP, NMF – non-negative matrix factorization, ProBEM – basophil eosinophil mast cell progenitor, ProgB – B cell progenitor, UMAP – uniform manifold approximation and projection.

**Figure S1:**
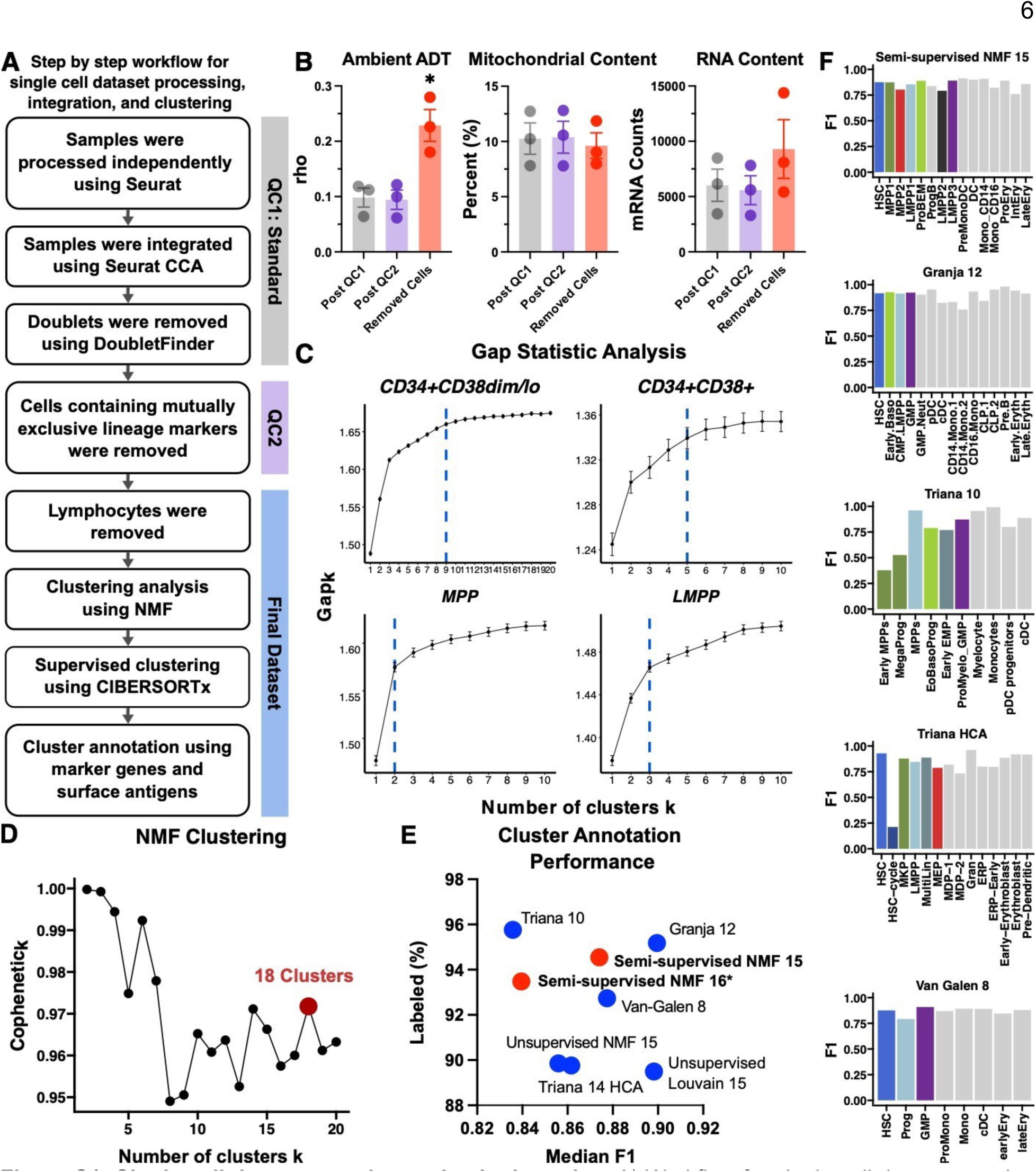
Single cell data processing and sub-clustering. A) Workflow for single cell data processing and clustering analysis. B) Ambient ADT signal (rho) of single cells after specific stages of data processing. C) Gap statistic analysis of cells in each respective immunophenotypic gate. D) Cophenetic indices of across k=2-20 clusters derived using NMF. E) Cell annotation transfer performance metrics comparing our new reference profile containing 15 clusters against published annotations and unsupervised clustering using the NMF algorithm (k=16; one cluster removed due to low cell content) and Louvain algorithm. F) Median F1 scores by cluster determined by intra-dataset annotation transfer (Methods) for select references. Colored bars highlight primitive HSPC clusters. ADT – Antibody derived tags, DC – dendritic cell, EMPP – erythroid-biased MPP, GMP – granulocyte monocyte progenitor, HSC – hematopoietic stem cell, HSPC – hematopoietic stem and progenitor cell, LMPP – lymphoid-primed MPP, MPP – multipotent progenitor, non-negative matrix factorization – NMF, ProBEM – basophil eosinophil mast cell progenitor, ProgB – B cell progenitor.

**Figure S2:**
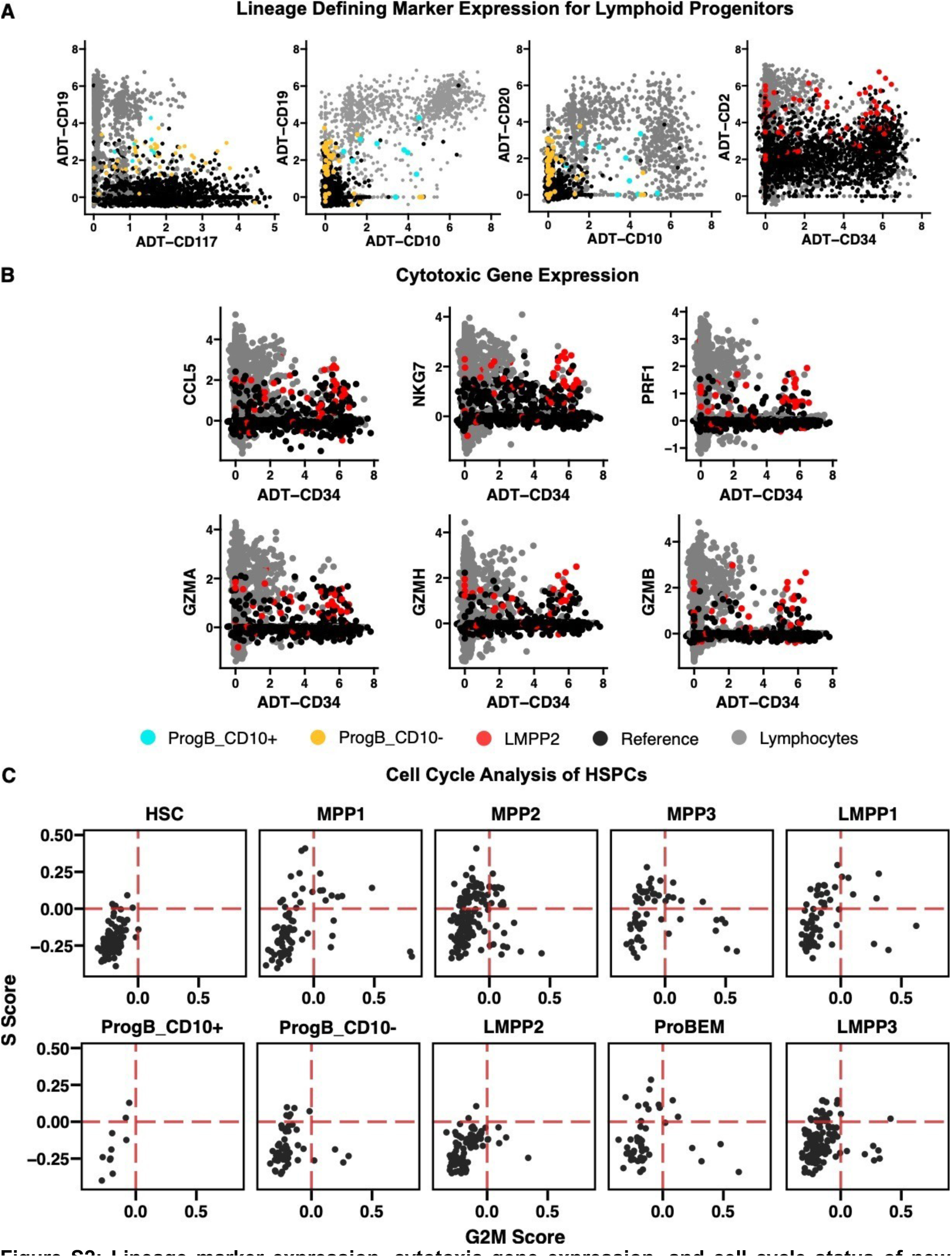
Lineage marker expression, cytotoxic gene expression, and cell cycle status of new HSPC sub-populations. A) HSPC cell cycle scoring. B) ADT plots illustrating key surface marker expression of certain HSPCs in relationship to remaining HSPCs (reference) and excluded lymphocytes. C) Comparison of cytotoxic gene expression between LMPP2 (red) and other celltypes. ADT – Antibody derived tags, DC – dendritic cell, EMPP – erythroid-biased MPP, GMP – granulocyte monocyte progenitor, HSC – hematopoietic stem cell, LMPP – lymphoid-primed MPP, MPP – multipotent progenitor, MPPP – myeloid-biased MPP, ProBEM – basophil eosinophil mast cell progenitor, ProgB – B cell progenitor.

### Clustering and computational purification of adult HSPCs

Prior scRNA-seq studies suggest that the HSPC sub-population in adult bone marrow is difficult to sub-cluster^14^. As an exploratory analysis of our new dataset, we first estimated the number of clusters within canonical HSPC sub-populations using both gene and surface marker expression. Single cells were computationally isolated using ADT counts and conventional markers used in FACS based gating (Table S4)^29^. The number of sub-clusters for each computationally purified sub-population was estimated using the Gap statistic^37^ on integrated features from concurrent whole transcriptome analysis (WTA) and ADT data (Figure S1C). Unlike prior work, we observed stable sub-clustering within CD34+CD38dim/lo sub-population (Figure S1C)^5^, which motivated an unsupervised sub-clustering analysis. We evaluated two clustering methods, the Louvain algorithm^38^ and non-negative matrix factorization (NMF)^39^, and observed that NMF based clustering generated more sub-clusters within the HSPC compartment (data not shown). We therefore performed NMF using the top 2000 features which revealed several optimal subclustering levels based on the cophenetic index (Figure S1D)^39,40^. To identify the optimal level of sub-clusters within the CD34+CD38dim/lo sub-population, we first split the data into a training and test dataset (Methods), and generated a signature matrix using the training dataset for deconvolution with CIBERSORTx^41^. The test dataset was then over-clustered (*k*=18; Figure S1D), and transcriptionally similar clusters were collapsed to optimize the deconvolution performance with CIBERSORTx in pseudo-bulk samples. This semi-supervised approach resulted in 15 high quality clusters (Figure 1B-D, Methods). We subsequently evaluated the strength of our subclustering approach by performing intra-dataset cell-annotation transfer (Figure S4, Methods). This reference profile (i.e. semi-supervised NMF 15) performed similar if not better than alternative clustering approaches (*k*=15 NMF and *k*=15 Louvain clustering) and published annotations^30,31,34^ based on bulk and cluster specific F1 scores (Figure S1E-F). Of note, our clustering approach allowed HSPC sub-population discrimination, whereas published annotations and the unsupervised Louvain approach generally combined these cells into a single cluster (Figure S1F). The final cluster annotations were initially derived based on marker gene and protein expression and include 4 sub-clusters within both the traditional MPP and LMPP subpopulations. Differential markers were used for the preliminary annotations (Figure 1D-F, Table S6), which includes 4 clusters within the canonical MPP gate: MPP1, MPP2, MPP3, and a basophil, eosinophil, and mast cell progenitor (ProBEM, annotated based on gene expression pattern). We also identified 4 clusters within the canonical LMPP gate: LMPP1, LMPP2, ProgB (annotated based on gene and surface marker expression, see below), and LMPP3.

Differentially expressed ADT counts were calculated iteratively to determine a new sorting strategy for these clusters (Figure 2A, Table S4, Methods). Computational purification using the new sorting strategy revealed greater target cluster enrichment and lower sub-population diversity compared to the traditional sorting strategy (Figure 2B-D). Improvements in purity were most noticeable for cells within the traditional MPP sub-population (Figure 2D). From our analysis, we identified CD69, CLL1, and CD2 as important surface antigens for purifying newly identified HSPCs, in addition to conventional markers CD34, CD38, CD45RA, and CD90. CD69 expression distinguished MPP1 and MPP2, and CLL1 expression marked MPP3 and LMPP3. We observed a candidate B cell progenitor, ProgB, in the canonical LMPP population, based on both gene and surface marker expression (Figure 2A, S2A-B, Table S6). Prior work has revealed that CD117, CD10, and CD19 are important markers for discriminating B cell progenitors^42^, so we evaluated ProgB cells for CD117, CD10, and CD19 expression (Figure S2A), and sub-annotated based on CD10 expression (Figure 1D). We found that the majority of ProgB cells express low levels of CD117 and CD19 but do not express CD10 (ProgB_CD10-), and only a rare population of ProgB cells express CD10 (ProgB_CD10+). Of note, both CD19 and CD10 expression was lower in ProgB than in lymphocytes. We also observed that CD2 expression marked a new candidate LMPP, LMPP2, which expresses cytotoxic genes at higher levels compared to other CD34+ HSPCs (Figure S2B). The complete immunophenotypes are reported in Table S4. Lineage defining marker expression compared to all cell types, including lymphocytes, are presented in Figure S2A, and cell cycle scores are presented in Figure S2C. As expected, HSCs exhibit low S and G2M scores, whereas the remaining HSPCs are upregulating both S and G2M signatures suggesting cell cycle entry at variable levels. Overall, our observations support a semi-supervised approach, i.e., unsupervised NMF clustering followed by manual curation with CIBERSORTx (Figure 1B-D), for identifying novel cell types in our single cell dataset.

**Figure 2:**
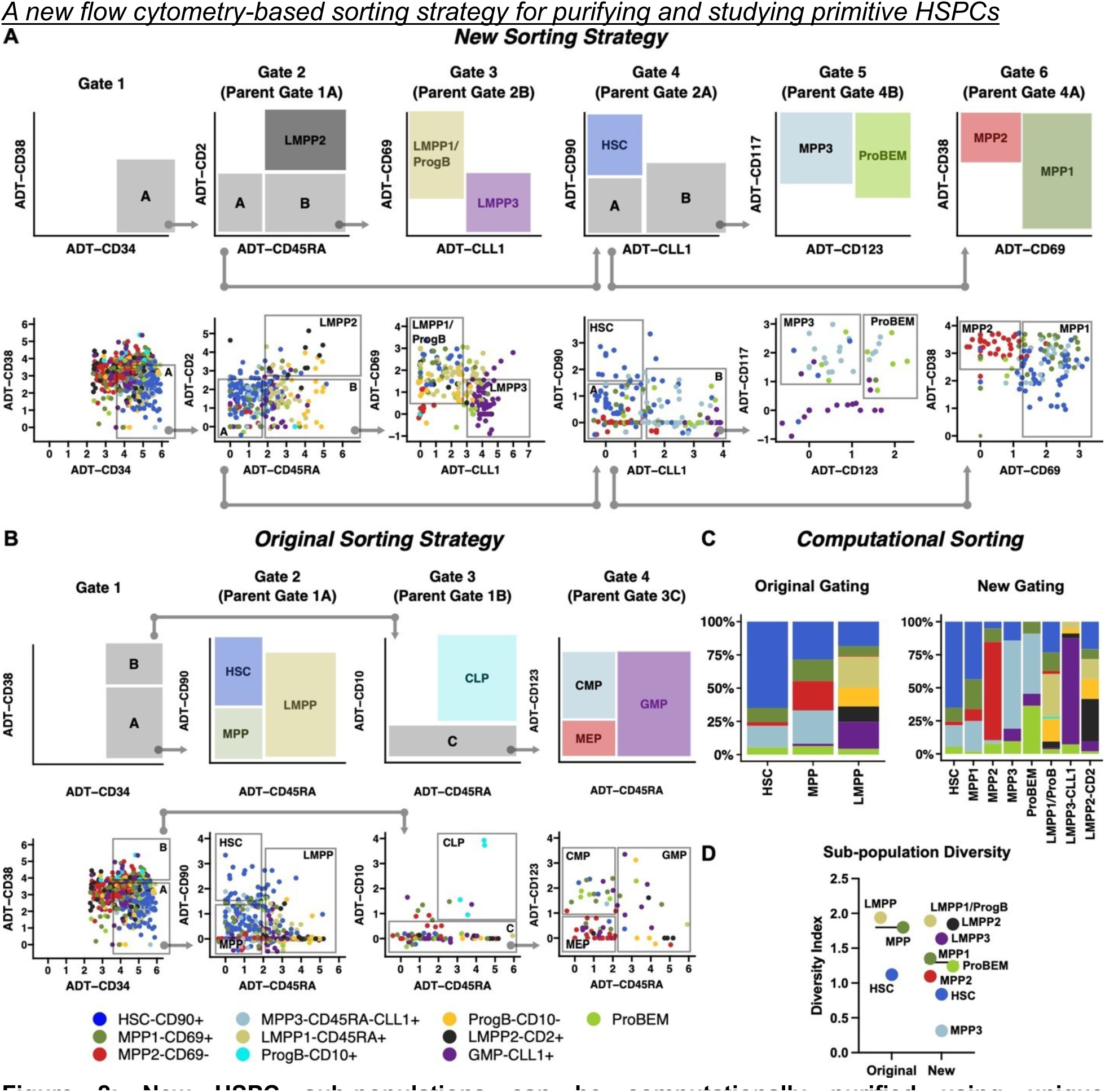
New HSPC sub-populations can be computationally purified using unique immunophenotypes. A) A schematic illustrating a new sorting strategy for purifying newly identified HSPC sub-populations. B) Schematic illustrating the original sorting strategy used for purifying HSCs, MPPs, and OPPs. C) Stacked bar plots illustrating cluster frequencies in target gates after computationally sorting cells using the original and new sorting strategy. D) Shannon diversity indices of target gates after computationally purifying cells using original and new sorting strategies (see Table S4). ADT – Antibody derived tags, DC – dendritic cell, EMPP – erythroid-biased MPP, GMP – granulocyte monocyte progenitor, HSC – hematopoietic stem cell, HSPC – hematopoietic stem and progenitor cell, LMPP – lymphoid-primed MPP, MPP – multipotent progenitor, MPPP – myeloid-biased MPP, oligopotent progenitor – OPP, ProBEM – basophil eosinophil mast cell progenitor, ProgB – B cell progenitor.

**Figure S3:**
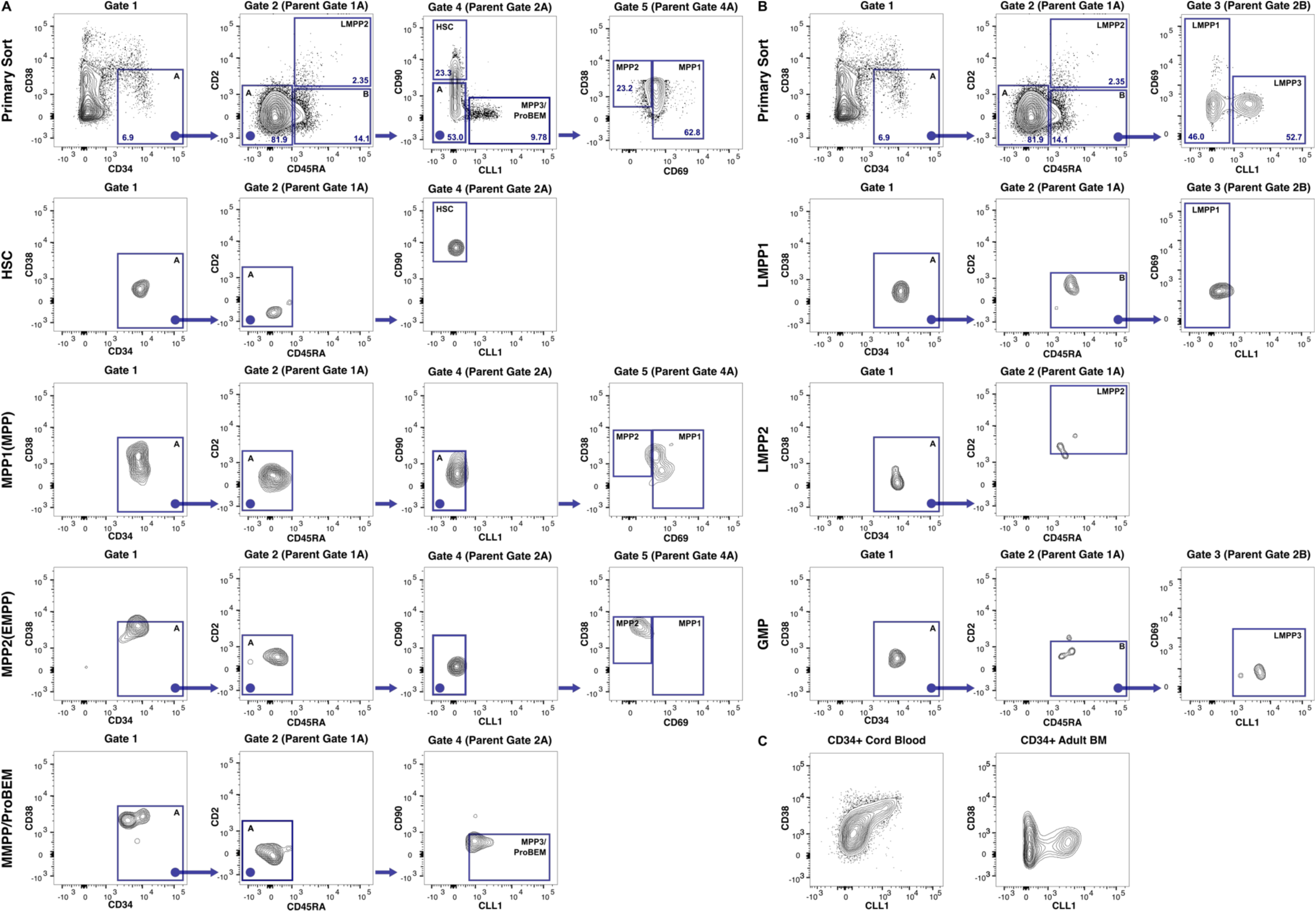
Representative plots of primary and post sorts of lineage negative adult bone marrow and cord blood cells. A) Sorting strategy for HSCs, MPP1 (MPPs), MPP2 (EMPPs), and MPP3 (MMPP)/ProBEMs with representative post sort purities. B) Sorting strategy for LMPP1-2 and LMPP3 (GMPs) with representative post sort purities below. C) Representative CD38 and CLL1 expression of CD34+ cells from adult BM or CB. BM – bone marrow, CB – cord blood, EMPP – erythroid-biased MPP, GMP – granulocyte monocyte progenitor, HSC – hematopoietic stem cell, LMPP – lymphoid-primed MPP, MPP – multipotent progenitor, MPPP – myeloid-biased MPP, ProBEM – basophil eosinophil mast cell progenitor.

### A new flow cytometry-based sorting strategy for purifying and studying primitive HSPCs

As the primary goal of this study is to understand the functional characteristics of our candidate subpopulations of MPPs and LMPPs, we designed a FACS strategy (Figure S3A-B) using the computationally derived immunophenotypes (Figure 2A, Table S4). Due to sort complexity and low cell numbers, MPP3s and ProBEMs were purified as one population. Additionally, CD19 was included as a lineage marker (Table S4), thereby excluding ProgB cells in these assays. We first used this final sorting strategy to evaluate the proportion of each cell type within the LinCD34+CD38dim/lo subpopulation across 24 adult BM and 4 CB donors (Figure 3A, Table S4) to compare the cellular composition between different donor sources. We observed a significant increase in MPP3/ProBEMs and LMPP3s and a decrease in MPP2s in adult bone marrow compared to CB (Figure 3A). The greater degree of heterogeneity in adult BM is partly marked by the expression CLL1 in CD34+CD38dim/lo cells compared to CB (Figure S3C). Prior work has demonstrated the utility of CD110, CD71, and CD49f in identifying HSC and MPP subsets in CB^13,43^. We therefore evaluated the expression of these markers among the newly identified HSPCs and observed greater CD71 expressing cells and fewer CD110 expressing cells in MPP2s compared to HSCs, MPP1s, and MPP3s (Figure 3B-E). Most HSCs and MPP1s expressed higher CD110 compared to MPP2 and MPP3s. There was no significant difference between the frequency of CD49f expressing cells within HSCs and new MPP cell types in either adult BM or CB (Figure 3F).

**Figure 3:**
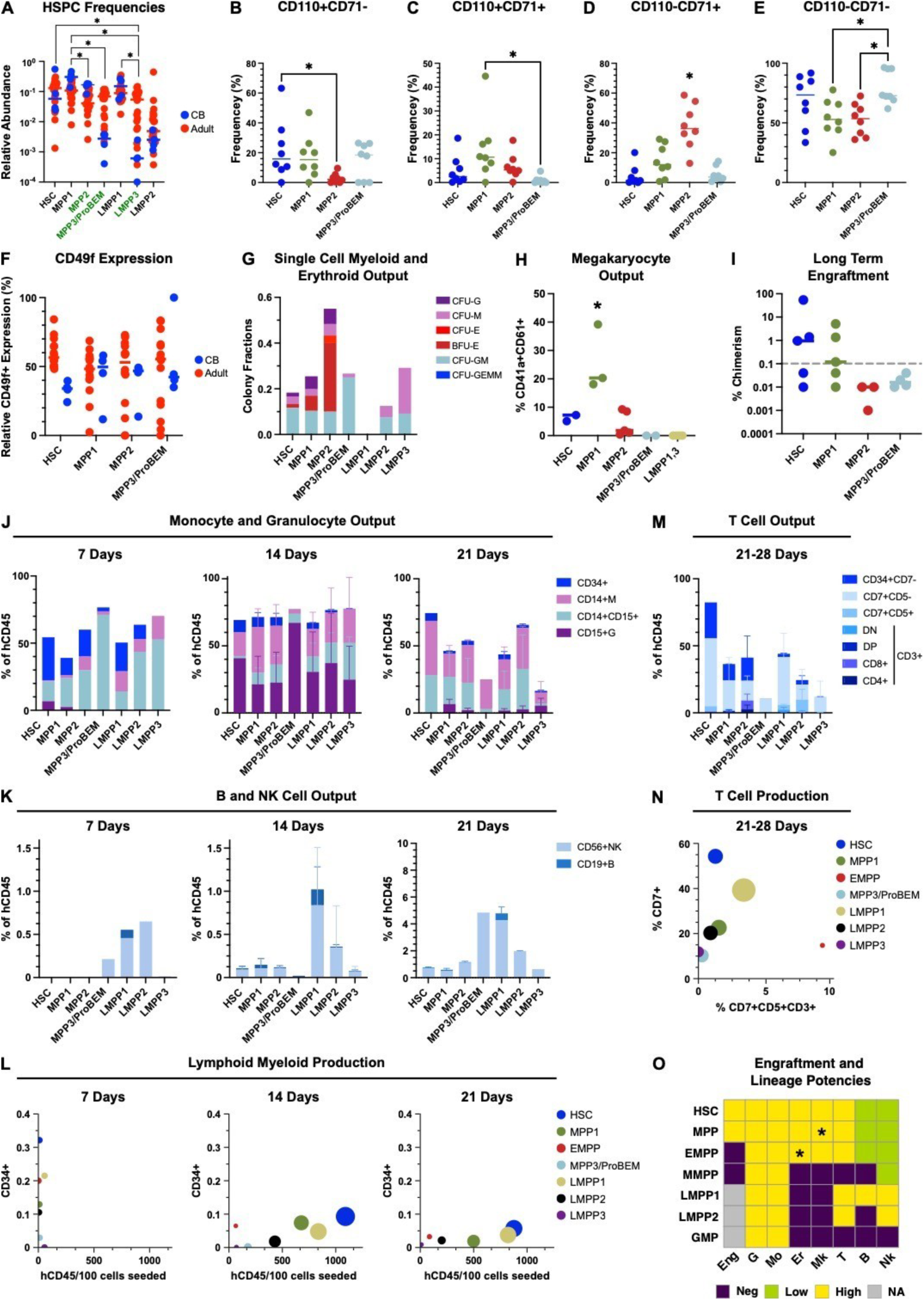
Computationally identified HSPCs can be purified using flow cytometry and exhibit distinct functional properties. A) HSPC frequencies across 24 adult bone marrow and 4 cord blood donors. B-E) Cells within each respective gate were evaluated for CD71 and CD110 expression and relative frequencies are displayed. F) CD49f expression of cells within each respective gate. G) Single cell methylcellulose colony-forming unit (CFU) assay of clonogenic output and lineage differentiation potencies of adult HSPCs. H) Proportions of CD41a+CD61+ megakaryocyte progenitors produced after 7-day in vitro expansion of each adult HSPC. I) Human to mouse chimerism 17-18 weeks post primary transplantation of purified HSPCs. Chimerism > 0.1% was considered positive for long term engraftment. Monocyte and granulocyte (J) and B and NK cell (K) output after 7-21 days of expansion in liquid culture with OP9 cells and SGF15 media. L) Lymphoid and myeloid production after 7-21 days of expansion. The size of each circle correlates with hCD45+ cell production normalized to 100 cells seeded. M) T cell output after 21-28 days of expansion in liquid culture with OP9-hDL4 cells and SF7 media. CD7+CD5+CD3+ cells were split based on CD4 and CD8 expression, and the remaining cells did not express CD3. N) T cell production after 21-28 days of expansion. The size of each circle correlates with hCD45+/SSAlo cell production normalized to 100 cells seeded. O) Summary of engraftment and lineage differentiation potencies for all HSPCs. DC – dendritic cell, DN – double negative, DP – double positive, E – erythroid, EMPP – erythroid-biased MPP, G – granulocyte, GEMM – granulocyte, erythrocyte, monocyte, megakaryocyte, GM – granulocyte monocyte, GMP – granulocyte monocyte progenitor, HSC – hematopoietic stem cell, HSPC – hematopoietic stem and progenitor cell, LMPP – lymphoid-primed MPP, MPP – multipotent progenitor, MPPP – myeloid-biased MPP, ProBEM – basophil eosinophil mast cell progenitor. *Adjusted p-val < 0.05.

Although candidate HSPCs were rare in adult BM, we were able to purify these subpopulations using the new sorting strategy (Figure S3A-B) for *in vitro* and *in vivo* assays. In single cell methylcellulose-based colony-forming-unit (CFU) assays, HSCs, MPP1s, and MPP2s had both myeloid and erythroid clonogenic potential, whereas MPP3s, LMPP2s, and LMPP3s had only granulocyte and monocyte potential (Figure 3G). MPP2s demonstrated the highest clonogenic output whereas LMPP1s did not expand in this assay. To evaluate megakaryocyte production, we assayed purified HSPCs (770-4000 cells/cell type/well; Table S7) *in vitro* in a serum-free liquid culture system. After 7 days of culture, MPP1s produced the greatest relative percentage of CD41a+CD61+ megakaryocyte progenitors (Figure 3H). HSCs and MPP2s produced minimal CD41a+CD61+ megakaryocyte progenitors whereas MPP3s and LMPP1s and LMPP3s did not produce megakaryocyte progenitors.

HSPC populations were subsequently evaluated for long-term engraftment in NOD/SCID/IL2Rγ/-(NSG) mice. Purified HSC and candidate MPPs (370-2970 cells/cell type/mouse; Table S8) were injected intrafemorally, and human chimerism was measured between 17-18 weeks^44^. HSCs and MPP1s were able to produce long-term engraftment compared to MPP2s and MPP3/ProBEMs (Figure 3I). In all engrafted mice, we observed the presence of immunophenotypically defined HSCs, MPP1s, MPP2s, and MPP3/ProBEMs, and both CD33+ and CD19+ cells within the human CD45+HLA+ sub-population (data not shown).

To characterize lineage potencies across all HSPCs, we performed a series of *in vitro* functional assays. Combined lymphoid and myeloid differentiation potential was assessed in liquid culture on OP9 cells with SCF, G-CSF, FLT3L, IL-2, DuP-697, and IL-15^6^. Purified HSPCs (200-2600 cells/cell type/well; Table S7) were cultured for 3 weeks and aggregate cellular output was quantified in Figure 3J-L. All cell types produced hCD45+CD14+ monocyte-macrophages (Ms) and CD15+ neutrophil-granulocytes (Gs), with a noticeable burst of Gs at 14 days (Figure 3J). LMPP1-2 cells produced the greatest amount of hCD45+CD14-CD15-SSAlowCD56+ NK cells, whereas HSCs, MPP1s, MPP2s, and MPP3/ProBEMs produced moderate to minimal NK cells (Figure 3K). LMPP1 cells also produced the greatest amount of hCD45+CD14-CD15SSAlowCD19+ B cells, whereas HSCs, MPP1s, and MPP2s demonstrated moderate to minimal B cell production (Figure 3K). Total cellular production was quantified throughout the 3 weeks, and HSCs, LMPP1, MPP1s, and LMPP2s had the greatest cellular output in this assay (Figure 3L).

T cell potency was assessed using an *in vitro* liquid culture system with OP9 stromal cells expressing human notch receptor ligand DL4 (OP9-hDL4)^45^. Purified HSPCs (200-2600 cells/cell type/well; Table S7) were cultured with cytokines SCF, FLT3L, and IL-7 for 21-28 days ^6^. HSCs, MPP1s, MPP2s, LMPP1, and LMPP2s produced moderate hCD45+SSAlowCD7+CD5+CD3+ cells with a few samples producing minimal CD4+, CD8+, or CD4+CD8+ DP cells (Figure 3M). MPP3s produced very few immunophenotypic T cells and LMPP3s produced no immunophenotypic T cells. LMPP1 cells generated the greatest total cellular production after 2128 days in culture (Figure 3N).

Both HSCs and MPP1s demonstrated long-term engraftment capacity and 7 lineage differentiation potential. MPP1s demonstrated greater megakaryocyte differentiation potential whereas MPP2s demonstrated greater erythroid potential compared to other HSPCs *in vitro*. Although, LMPP1s had the greatest lymphoid potential *in vitro*, we note that all HSPCs evaluated in our assays except for LMPP3s contain a degree of lymphoid potency. Based on the results of the functional assays, MPP1 was annotated as an MPP, MPP2 was annotated as an erythroidprimed MPP (EMPP), MPP3 was annotated as a myeloid-primed MPP (MMPP), and LMPP3 was annotated as a GMP. Long-term engraftment capacity and 7 lineage differentiation potencies are summarized for each HSPC population in Figure 3O.

### An updated model of adult human hematopoiesis

An updated model of adult hematopoiesis based on our complementary computational and functional analyses is presented in Figure 4. Each HSPC’s differentiation potential is designated by arrows with thickness corresponding to the strength of the lineage potency. Annotations and arrows that were not directly studied in our functional assays are marked with dashed lines and were incorporated into the model based on our computational analysis and prior research observations. The canonical CD90+ HSC can be further purified by removing CD2+ and CLL1+ cells. The 4 clusters within the canonical CD90-CD45RA-MPP gate can be purified using CD69 and CLL1 and show distinct functional properties (Figure 3). The canonical CD90CD45RA+LMPP contain 4 additional clusters as well that can be purified using CD2, CLL1, and CD69. ProgBs and ProBEMs could be purified using CD123, CD117, CD19, and CD10, but this was not evaluated in the current study.

**Figure 4:**
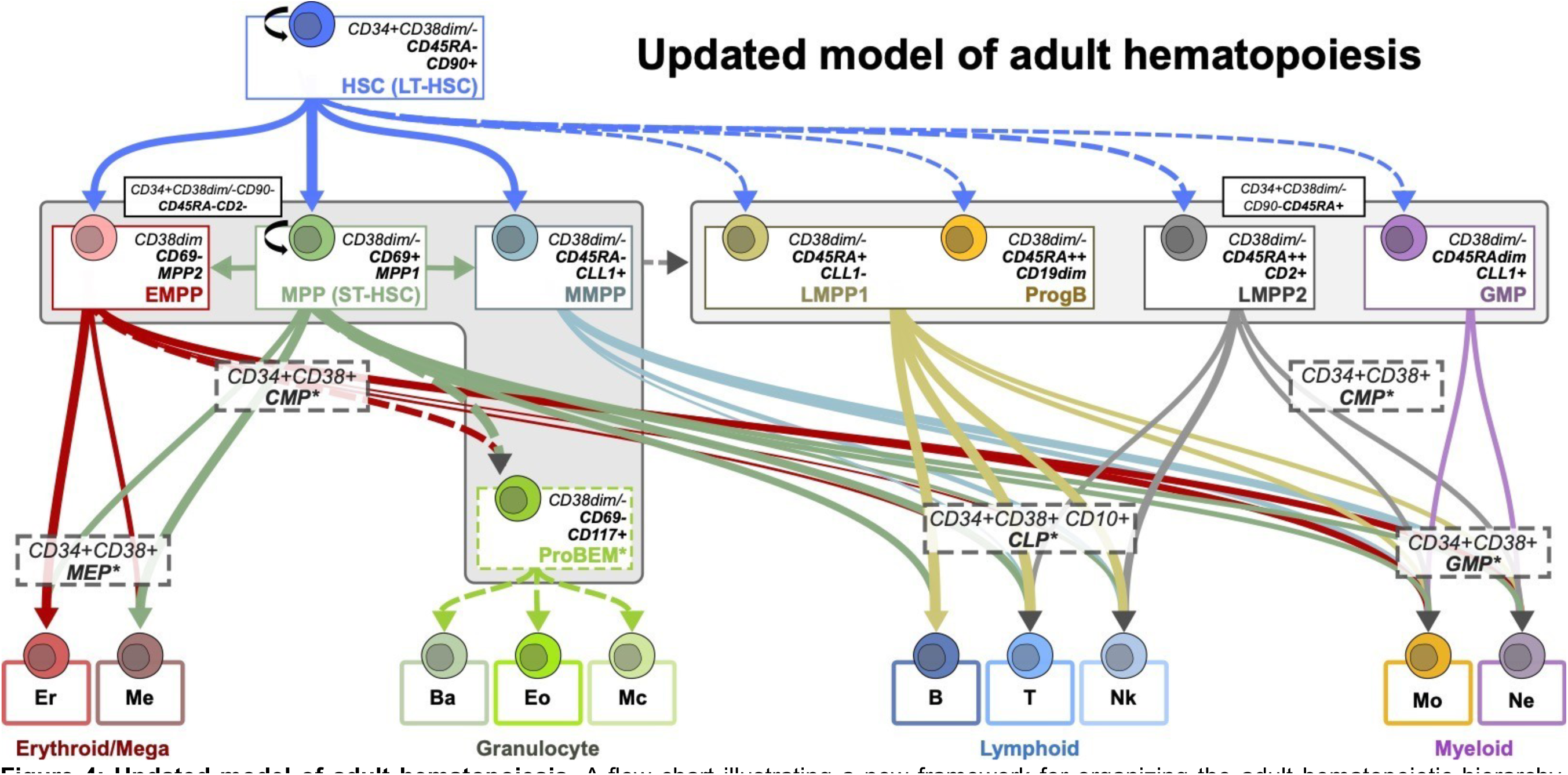
Updated model of adult hematopoiesis. A flow chart illustrating a new framework for organizing the adult hematopoietic hierarchy. Arrows depict differentiation potencies with relative thickness highlighting the strength of that potency. Dashed lines designate cell populations and differentiation trajectories that have been previously described but not evaluated in this study. The updated model highlights the heterogeneity within the canonical MPP and LMPP populations in adult bone marrow. We propose that within the canonical Lin-CD34+CD38dim/loCD90-CD45RA-MPP sub-population, the MPP resides in CLL1CD69+ cells, and CLL1+ cells are myeloid-biased whereas CLL1-CD69-cells are erythroid-biased. Within the canonical Lin-CD34+CD38dim/loCD90-CD45RA+ LMPP population, we propose that CD2-CLL1-LMPP1s have the strongest lymphoid potency, CD2+ LMPP2s have both lymphoid and myeloid potency with high expression of cytotoxic genes, and CD2-CLL1+ cells are primarily GMPs. Ba – basophils, CLP – common lymphoid progenitor, CMP – common myeloid progenitor, EMPP – erythroid-biased MPP, Eo – eosinophils, Er – erythrocytes, GMP – granulocyte monocyte progenitor, HSC – hematopoietic stem cell, LMPP – lymphoid-primed MPP, Mc – mast cells, Me – megakaryocytes, Mo – monocytes, MPP – multipotent progenitor, MPPP – myeloid-biased MPP, Ne – neutrophils, ProBEM – basophil eosinophil mast cell progenitor, ProgB – B cell progenitor.

### New MPP populations are associated with unique changes in TF networks

Transcription factors (TF) modulate specific gene regulatory networks (GRNs), which are critical for determining cell identity, function, and lineage commitment^15,29,46,47^. Studies evaluating TF GRNs in hematopoiesis suggest that specific cell fate and lineage decisions are regulated by complex circuits involving several TFs rather than the sequential expression of individual, lineage specific factors^48,49^. These observations may be particularly relevant for HSPCs which can be regulated by small variations in several unique TFs^47,49^. We therefore evaluated if our new HSPC cell states could be characterized by specific TFs or a combination of several factors. Motif analyses have been useful for evaluating differential GRNs and cell states^29^, however TF families often share similar DNA binding motifs which confounds the analysis. To address these questions and associated limitations, we used our new HSPC cell states as a reference profile to analyze published scATAC-seq data from healthy CD34+ adult BMMCs^30^ using the Analysis of Regulatory Chromatin in R (ArchR) framework (Methods)^50^. We were able to identify important TFs in adult hematopoiesis by utilizing an integrated gene expression and chromatin accessibility analysis and a non-redundant, multi-model motif reference database curated by Vierstra et al^51^. The integrated approach allowed us to discriminate between specific TFs within TF families, and the curated motif reference provided information across multiple models while retaining TF binding information. Specifically, scATAC-seq cells were integrated with scRNA-seq cells using inferred gene scores (scATAC-seq) with measured gene expression (scRNA-seq) (Figure 5A-B, Methods). This enabled cross platform cell annotation transfer and gene expression integration of scATAC-seq cells. The inferred gene expression matrix, ie gene integration matrix (GIM), for the annotated scATAC-seq cells are shown in Figure 5B and the expected reference gene expression (GEM) for the same genes are shown in Figure 5A. Averaged normalized counts per cluster for these top differentially expressed genes were highly correlated between the two gene expression matrices (Figure 5C). The integrated dataset was first used to evaluate correlated motif accessibility with TF gene expression (putative regulators). Given the underlying noise associated with cross platform cell annotation transfer and gene expression inference, we identified putative regulators that also varied significantly across differentiation trajectories based on our functional studies (differential regulators) to identify TFs that were most likely to influence HSPC GRNs (Figure 5D, Methods). As the new cell types observed from these studies reside in the canonical HSC and MPP sub-populations (HSC, MPP, EMPP, MMPP, and ProBEM) we focused the TF GRN analysis on these specific HSPCs. Representative footprint plots for important TFs identified using the differential regulator approach are provided in Figure 5E, and a summary of the results are illustrated in Figure 5F. We observe that *TGIF2*, *REL*, and *ETS1* motifs are more accessible in HSCs compared to MPPs and MMPPs, and *ETV6*, *FLI1*, *ERG*, and *TCF4* motifs are more accessible in MPPs compared to HSCs, EMPPs, and ProBEMs. We also see that TF accessibility for primitive HSC/MPPs is lower than more differentiated progenitors, ie *TAL1* in EMPPs (Figure 5E). Our integrated analysis suggests that a network of lineage-defining TFs gain activity in primitive progenitors based on chromatin accessibility associated with specific HSPC cell states.

**Figure 5:**
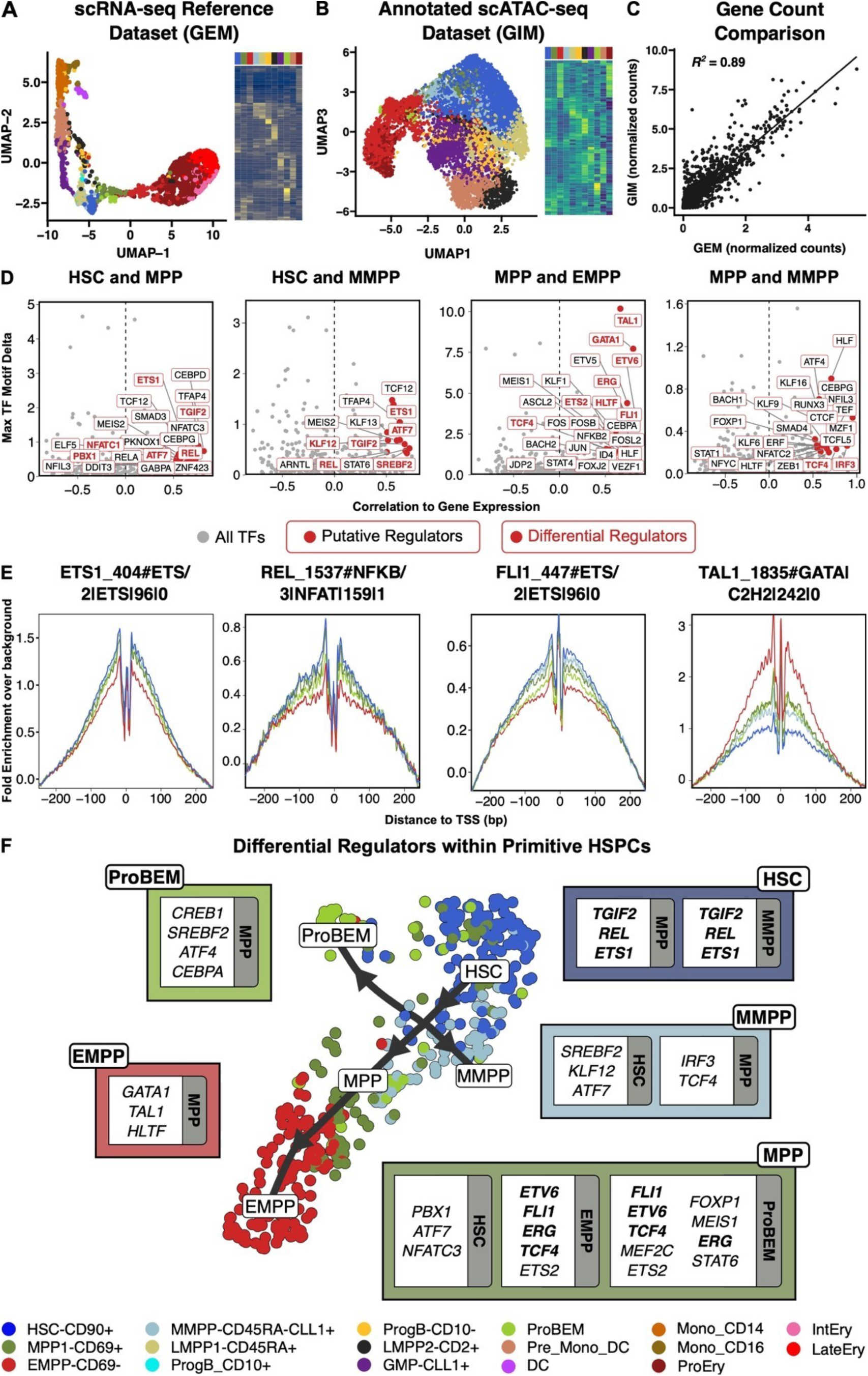
New adult HSPCs express distinct changes in chromatin accessibility. A) UMAP of scRNAseq reference and heatmap of top differentially expressed genes. B) UMAP projection of scATAC-seq cells labeled based on integration with scRNA-seq data and heatmap of the differentially expressed genes within the gene integration matrix. C) Correlation between averaged normalized counts per cluster for top differentially expressed features in the GIM (scATAC-seq dataset) and GEM (scRNA-seq reference dataset). D) Putative and differential regulators between designated HSPCs. Positive values are up in the first celltype indicated in the title. D) Footprints for representative differential regulators. E) Summary of key differential regulators that define each specific HSPC. Each box contains significant regulators compared to the labeled HSPC. Entries in bold are present in multiple comparisons. DC – dendritic cell, EMPP – erythroid-biased MPP, GMP – granulocyte monocyte progenitor, HSC – hematopoietic stem cell, LMPP – lymphoid-primed MPP, MPP – multipotent progenitor, MPPP – myeloid-biased MPP, NMF – non-negative matrix factorization, ProBEM – basophil eosinophil mast cell progenitor, ProgB – B cell progenitor.

**Figure S4:**
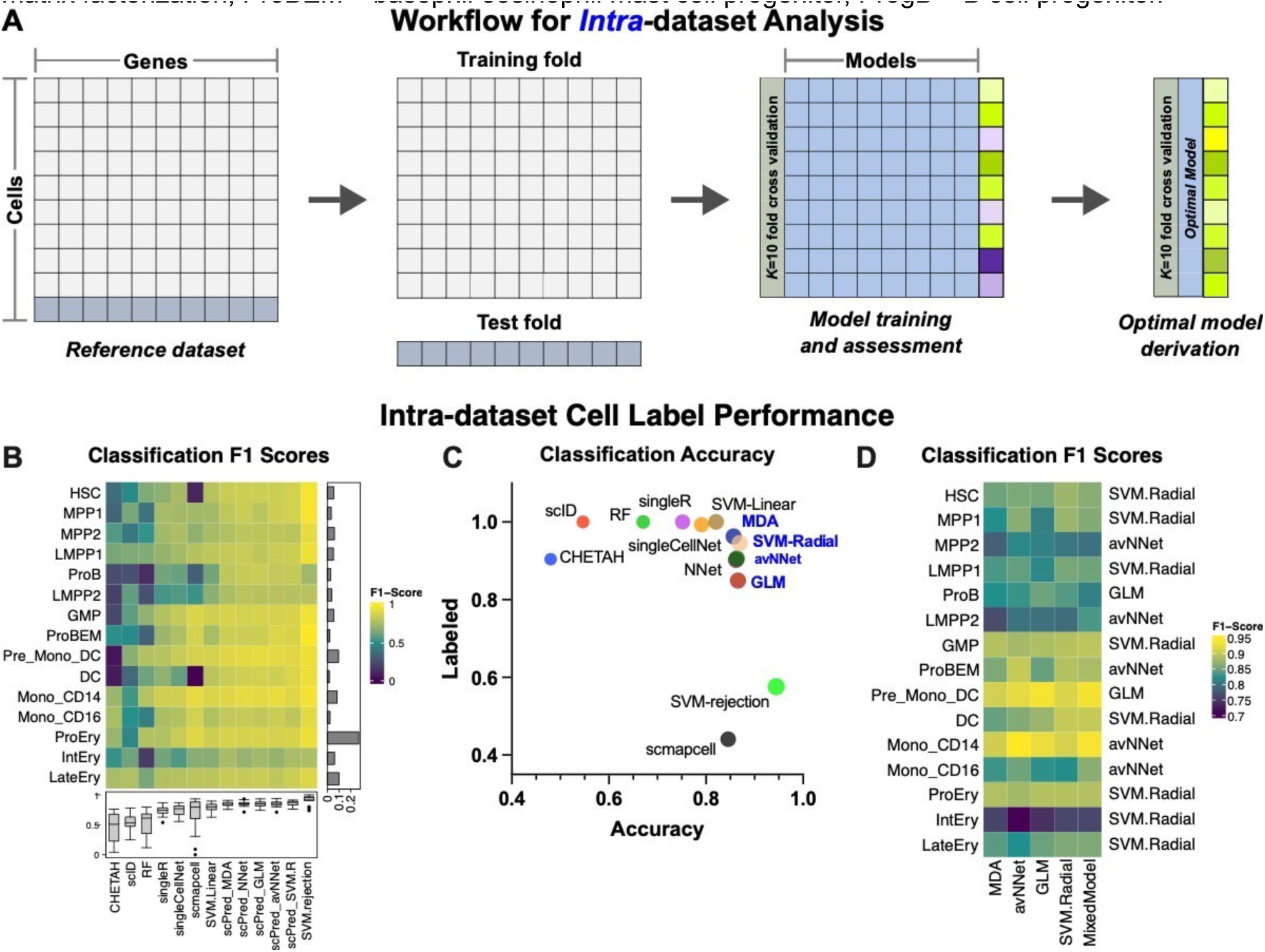
A mixed-model classifier accurately identifies similar cell states based on intra-dataset cell annotation transfer. A) Workflow for intra-dataset analysis. B) Classification F1 scores for each mode. Boxplots show range of F1 scores for each classifier and barplots represent proportion of each celltype. C) Performance of each classifier measured by proportion labeled and accuracy (median F1-score). D) Celltype specific F1 scores using high performing classifiers. Top performing classifier is indicated on the right. DC – dendritic cell, EMPP – erythroid-biased MPP, GMP – granulocyte monocyte progenitor, HSC – hematopoietic stem cell, HSPC – hematopoietic stem and progenitor cell, LMPP – lymphoid-primed MPP, MPP – multipotent progenitor, MPPP – myeloid-biased MPP, NMF – non-negative matrix factorization, ProBEM – basophil eosinophil mast cell progenitor, ProgB – B cell progenitor.

**Figure S5:**
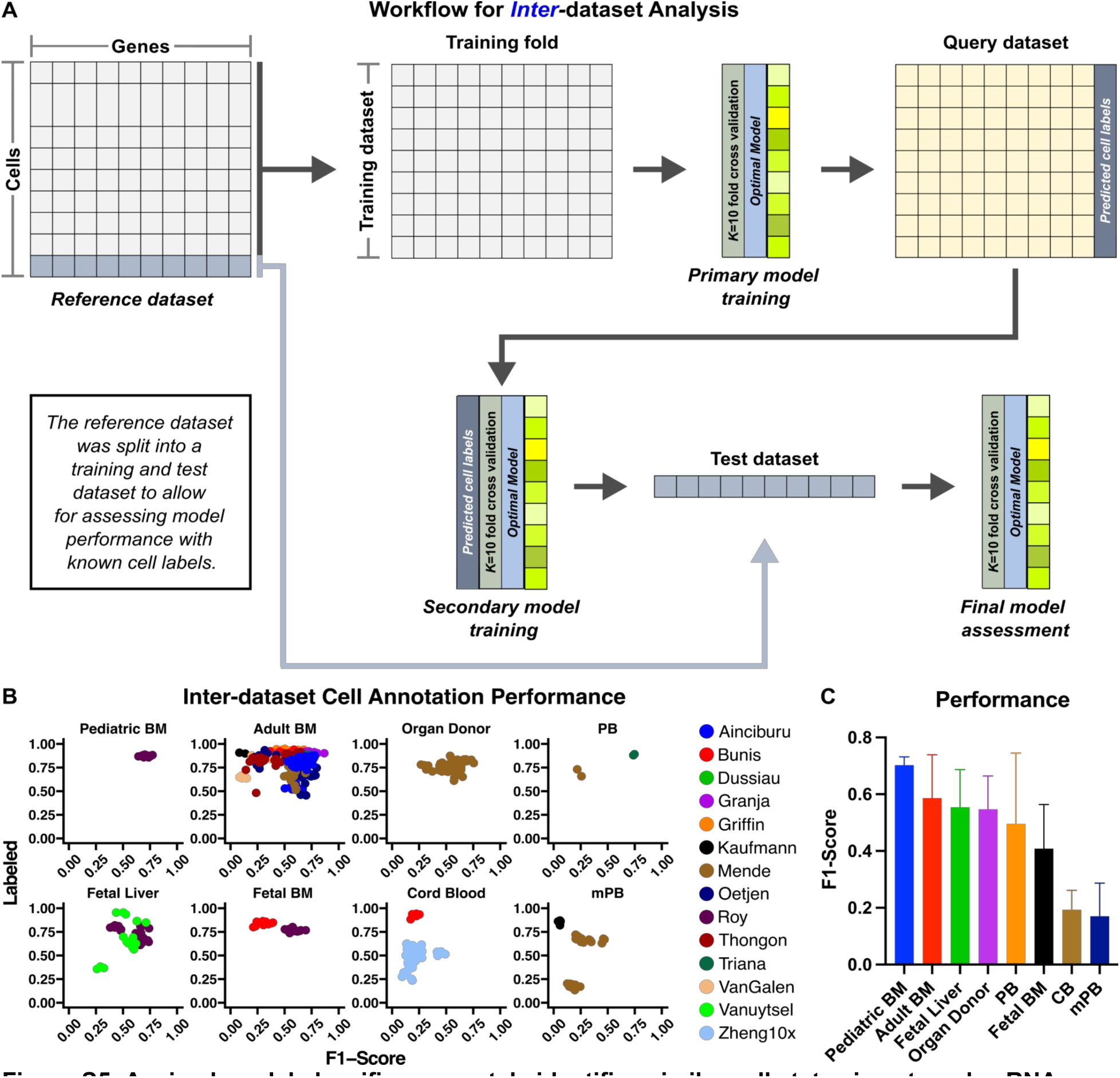
A mixed-model classifier accurately identifies similar cell states in external scRNA-seq datasets. A) Workflow for inter-dataset analysis. B) Performance of the mixed model classifier based on the inter-dataset analysis across all datasets. C) Mean F1 scores by donor source. BM – bone marrow, CB – cord blood, mPB – mobilized PB, PB – peripheral blood.

**Figure S6:**
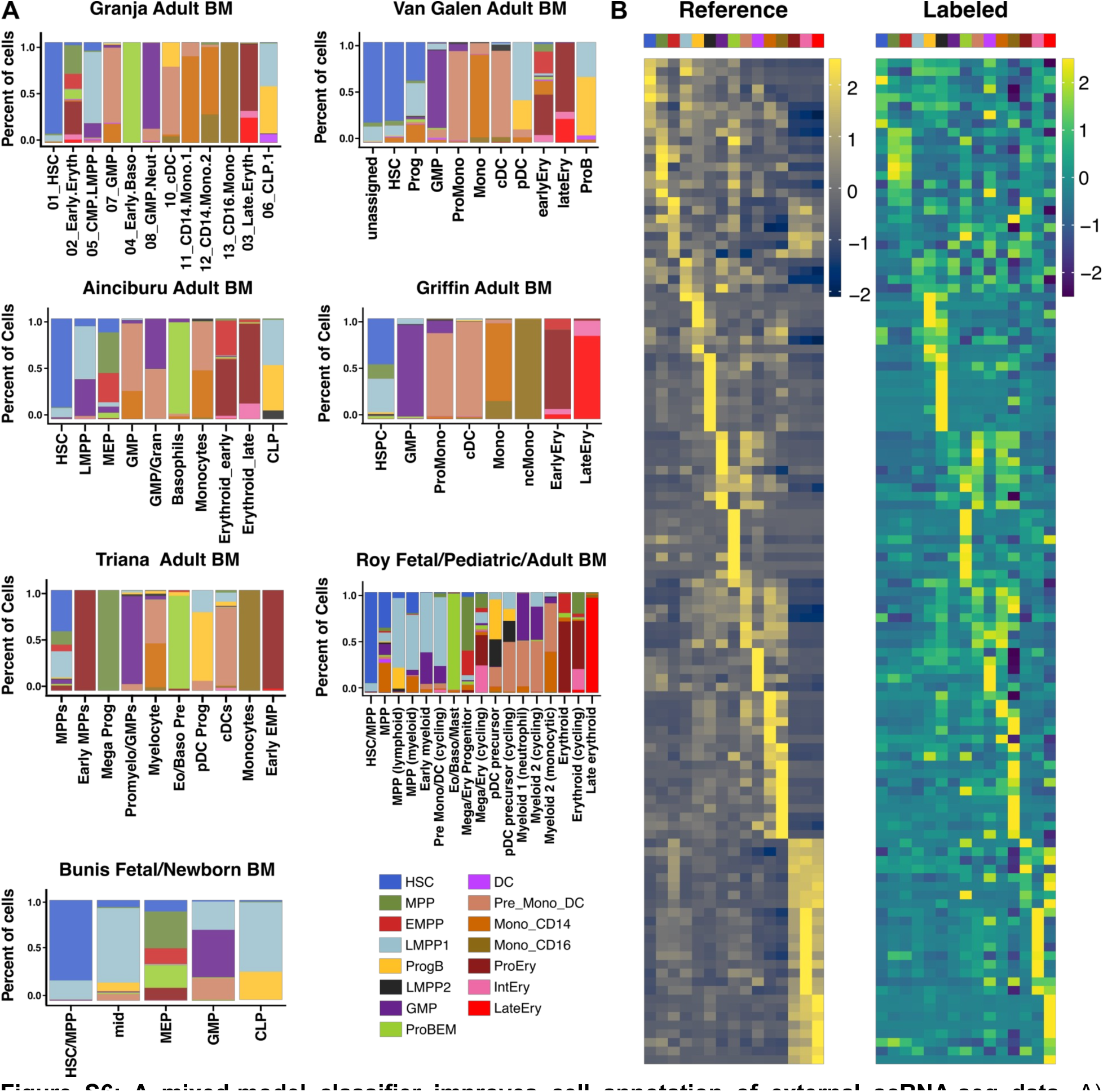
A mixed-model classifier improves cell annotation of external scRNA-seq data. A) Frequency of labeled cells within original annotations for evaluable datasets. B) Heatmap comparing expression of the top differentially expressed genes in the reference and labeled datasets. DC – dendritic cell, EMPP – erythroid-biased MPP, GMP – granulocyte monocyte progenitor, HSC – hematopoietic stem cell, LMPP – lymphoid-primed MPP, mobilized PB – mPB, MPP – multipotent progenitor, MPPP – myeloidbiased MPP, NMF – non-negative matrix factorization, peripheral blood – PB, ProBEM – basophil eosinophil mast cell progenitor, ProgB – B cell progenitor.

### A mixed model classifier improves cell annotation transfer across scRNA-seq datasets

After identifying functionally and molecularly distinct progenitors in adult BM, we tested if we could identify these populations in external scRNA-seq datasets. An important requirement for such an analysis is accurate cell annotation transfer using a supervised classification method. The performance of existing classification methods can be highly variable which in turn confounds interpretation of their cell annotation predictions. To identify the optimal cell classifier, we adapted a method developed by Abdelaal et al^52^ which now incorporates both model and cell type specific metrics (Figure S4-5). Classifier performance was evaluated using the 15 cell type low level annotation (Figure 1D, Table S6) and two metrics: F1 scores, which measures model accuracy, and cell annotation recovery (labeled), which quantifies the proportion of cells with confident cell annotation assignments. We first performed an intra-dataset analysis (Figure S4A) where we split our reference dataset into a test and training set and evaluated each model using 10-fold cross validation (CV). We evaluated 13 established classifiers (Figure S4B-C, Methods)^52^ using this approach, and found that high performing classifiers used support vector machines (SVMs), generalized linear models (GLMs), averaged neural networks (avNNets), or mixture discriminate analysis (MDA). We then evaluated the performance of each classifier for each cell type using the scPred R package^52^ which revealed that classifier performance was also dependent on cell type (Figure S4D). The optimal cell type specific method was incorporated into the final classifier, ie a mixed model classifier, and was evaluated further using an inter-dataset analysis (Figure S5A, Methods). We evaluated the performance of the mixed model classifier on 14 external scRNA-seq datasets^30,31,34,35,53–62^. The model performed well on both adult and pediatric bone marrow cells (Figure S5B); however, the performance was significantly reduced when annotating mobilized peripheral blood (mPB) and CB (Figure S5C). We also evaluated model performance by comparing our reference annotation assignments to the original cell annotations in 6 datasets containing this information and found good concordance (Figure S6A). Our annotation assignments allowed for further deconvolution of progenitor cell states in the external data.

Heatmaps illustrating marker gene expression for our reference dataset and the integrated external datasets are presented in Figure S6B. The optimized mixed model classifier was used for the subsequent analysis of external mouse and human scRNA-seq data.

### Human MPPs and OPPs show limited homology with mouse HSPCs

Research in murine hematopoiesis has identified immunophenotypically distinct HSPCs with unique functional characteristics^16,17^. These include sub-populations of murine HSCs, ie CD45+EPCR+CD48−CD150+ (ESLAM) HSCs, LT-HSCs, ST-HSCs^63,64^, and MPPs with varying lineage potencies, ie MPP2-4^16^ and MPP Ly I/II^65^. Delineating whether similar cell types exist in human adult hematopoiesis is critical for improving our understanding of human blood biology. Further, since the murine model is often used as a reference for human hematopoiesis, it will be important to understand homologous cell types between mice and humans to effectively use the mouse model as a reference. Because our analysis provided evidence that lineage biased MPPs exist in humans as well, we next attempted to identify their homologous cell states in murine hematopoiesis. To perform this analysis, we evaluated two single cell datasets with concurrent single cell mRNA and surface marker quantification in young adult (2-3 months)^64^ and adult (8-10 months) mice BM^65^. These data contained HSC and MPP annotations based on immunophenotype, which allowed for a cell type specific cross species analysis with our reference human HSPC profiles (Figure 6A). We first performed a gene ortholog-to-ortholog conversion of the mouse data and annotated the murine cells using our mixed model classifier (Figure 6B). In young adult mice, LT and ST-HSCs were all annotated as human HSCs, however, the ESLAM HSCs and MPP2-3 annotations were more heterogeneous. Murine MPP4s appear to resemble human LMPP1, murine MPP2s were mostly labeled as human MPPs and LMPP1s, and murine MPP3s were mostly labeled as LMPP1s. In adult mice, there was also a strong one-to-one association between HSCs. Murine granulocyte and monocyte progenitors (pGM_My) and CLPs were predominantly labeled as GMPs and LMPPs respectively, however, the assignments for the remaining cell types were more inconsistent (Figure 6B).

**Figure 6:**
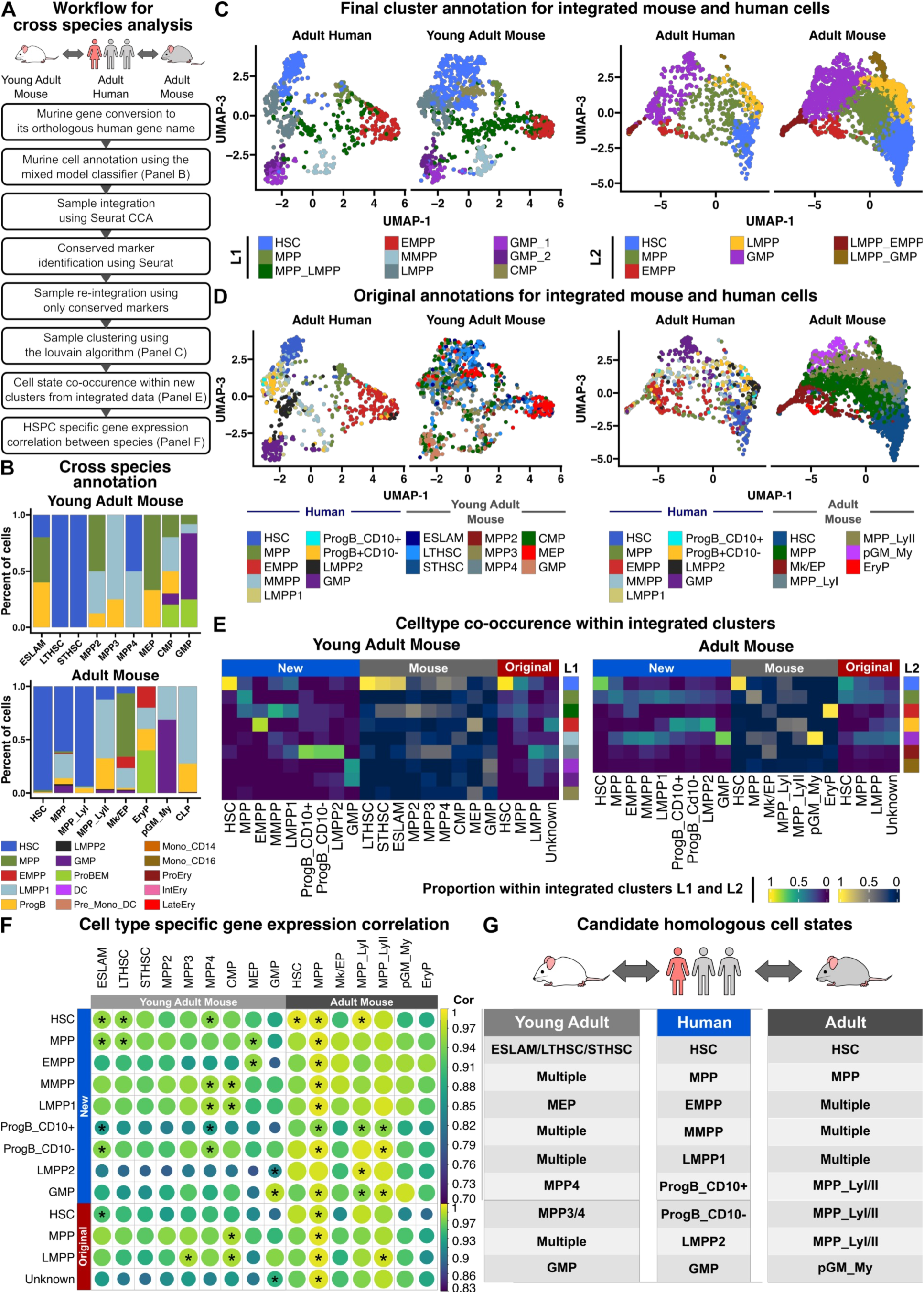
Human MPPs and OPPs have limited homology with corresponding murine HSPCs. A) Workflow for cross species analysis. B) Mouse cells were annotated with human HSPC labels using the mixed model classifier. Relative frequencies of each human celltype annotation are illustrated for each mouse HSPC. UMAP projections of integrated human and mouse cells colored based on C) new cluster labels L1 and L2 of integrated cells and D) original cell annotations for the mouse and human cells. E) Human and mouse HSPC co-occurrence plots. Each row is colored based on integrated cluster assignments L1 and L2 (see panel C), and each entry is colored based on the proportion (column comparison) of the original cluster within the new cluster assignments. F) Average gene expression correlation between human and mouse HSPCs. Correlation strength is reflected by both heat and circle size. *Relationship with highest correlation. G) Summary of candidate homologous states between human and mouse HSPCs. DC – dendritic cell, EMPP – erythroid-biased MPP, GMP – granulocyte monocyte progenitor, HSC – hematopoietic stem cell, HSPC – hematopoietic stem and progenitor cell, LMPP – lymphoid-primed MPP, MPP – multipotent progenitor, MPPP – myeloid-biased MPP, ProBEM – basophil eosinophil mast cell progenitor, ProgB – B cell progenitor.

**Figure S7:**
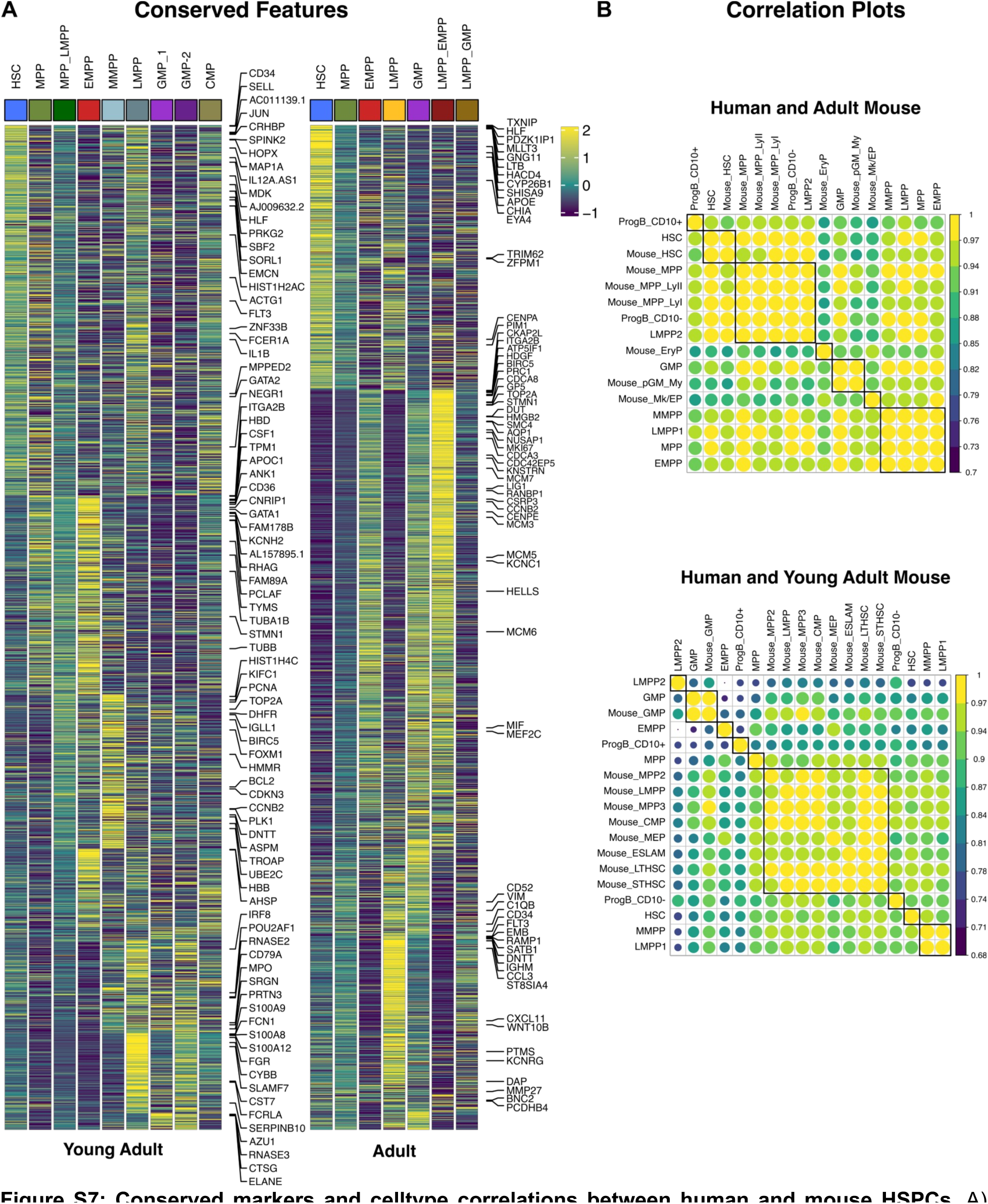
Conserved markers and celltype correlations between human and mouse HSPCs. A) Heatmap showing expression of conserved markers between human and mouse HSPCs. B) Hierarchal clustering celltypes based on average gene expression. EMPP – erythroid-biased MPP, GMP – granulocyte monocyte progenitor, HSC – hematopoietic stem cell, HSPC – hematopoietic stem and progenitor cell, LMPP – lymphoid-primed MPP, MPP – multipotent progenitor, MPPP – myeloid-biased MPP, NMF – nonnegative matrix factorization, ProBEM – basophil eosinophil mast cell progenitor, ProgB – B cell progenitor.

To improve our ability to identify homologous cell states, we performed an iterative integration of the murine and human data (Figure 6A, Methods). We first integrated each murine dataset with the human reference using Seurat CCA and identified conserved features between mouse and human HSPCs (Figure S7A, Table S11, Methods). The data were then re-integrated using only conserved features, and clustered based on the number of cell types present in the original murine dataset. The final clusters, L1 and L2, were annotated based on marker genes and their resemblance to our new human HSPC states (Figure 6C). For comparison, a separate projection displays the integrated cells based on the original species-specific annotation (Figure 6D). We then performed a cell type co-occurrence analysis (Figure 6E) by displaying the proportion (column comparison) of human HSPCs that co-occur in the same integrated clusters (L1 and L2) as murine HPSCs (row comparison). We evaluated human HSPCs based on our new sorting strategy (New, Figure 2A) as well as the original sorting strategy (Original, Figure 2B). We also compared average cell type specific gene expression correlation between human and mouse HSPCs using the same annotation framework (Figure 6F and Figure S7B). In young adult mice, we again saw homology between HSCs, and moderate homology between EMPPs and MEPs, ProgB and MPP4, and GMP states. In adult mice, we also saw homology between HSC states and between human GMPs and mouse pGM_My cells (Figure 6E). There was moderate homology between ProgB, LMPP2s and MPP_LyI/II cells. There were no clear associations between other cell types. A summary of candidate homologous cell states is listed in Figure 6G. These results indicate that while there are some similarities in the MPP subpopulations between mouse and human, there is not an exact correspondence in cell states indicating the potential for diversity among these cells in hematopoiesis.

### HSPCs undergo cell type specific changes in gene expression during aging

The long-term goal for developing a new classification framework for HSPCs is to improve our ability to study human health and disease. Aging is a primary risk factor for numerous diseases, and is often characterized by hallmarks including genomic instability, senescence, epigenetic changes, altered proteostasis, metabolic and mitochondrial dysfunction, maladaptive cellular signaling, and stem cell exhaustion^66^. In hematopoiesis, the relative contribution of these individual hallmarks to aging is unclear as is the interconnection between these processes and the associated cellular responses. We hypothesized that by performing a cell type specific analysis of human aging, we could overlay a cell specific context to classical derangements observed in human aging. To address this hypothesis, we evaluated whether our new HSPC cluster profiles (Figure 1D) can improve the ability to identify gene expression programs associated with human aging (Figure 7A). Unlike in mice, we cannot prospectively study aging HSPCs in humans. However, with the emergence numerous scRNA-seq datasets, we can indirectly evaluate aging with greater resolution using primary human data. To perform this analysis, we first identified six single cell datasets from human BM with age and cell type annotations^31,34,35,53,54,57^. Each dataset was labeled using the mixed model classifier and integrated in Seurat. After integration, we obtained 42,480 cells with high quality cell annotations across 20 donors aged 0-75 (Figure 7B).

**Figure 7:**
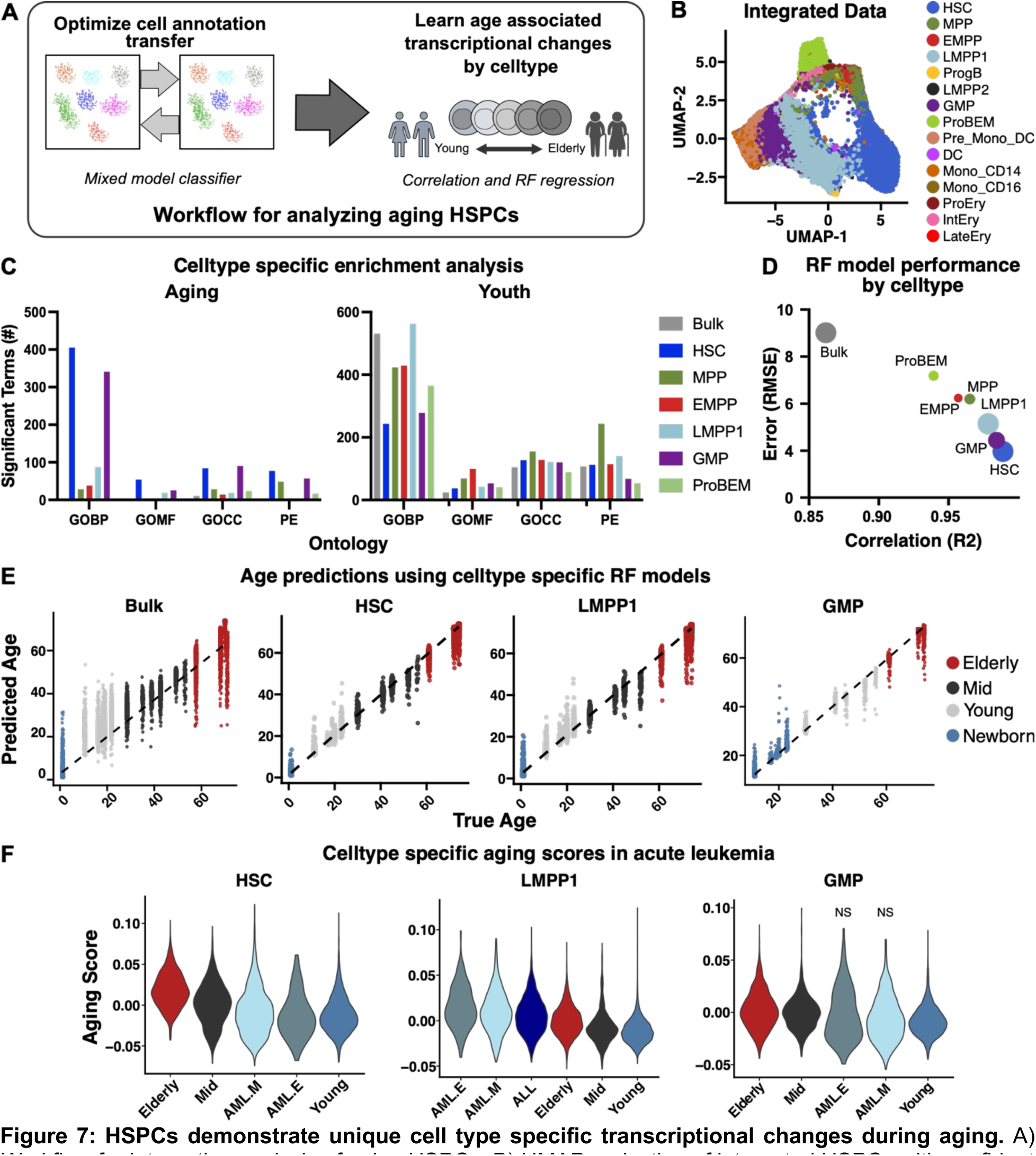
HSPCs demonstrate unique cell type specific transcriptional changes during aging. A) Workflow for integrative analysis of aging HSPCs. B) UMAP projection of integrated HSPCs with confident cell annotations. C) Enrichment analysis was performed using informative genes identified from celltype specific ELN regressions for aging and youth. Barplots show the number of significant terms for each celltype specific analysis. D) RF model performance metrics after RFE for predicting celltype specific age on the test data. E) Correlation between the predicted age from the RF model after RFE and true age on the test data. F) Celltype specific aging scores for healthy and cancer cells annotated using the mixed model classifier. Scores were significantly different unless marked NS. ALL – acute lymphoblastic leukemia, AML – acute myeloid leukemia, DC – dendritic cell, ELN – elastic net, EMPP – erythroid-biased MPP, GO – gene ontology, GOBP – GO biological processes, GOCC – GO cellular component, GOMF – GO molecular functions, GMP – granulocyte monocyte progenitor, HSC – hematopoietic stem cell, HSPC – hematopoietic stem and progenitor cell, LMPP – lymphoid-primed MPP, MPP – multipotent progenitor, MPPP – myeloid-biased MPP, NMF – non-negative matrix factorization, NS – not significant, PE – pathway enrichment (Reactome), ProBEM – basophil eosinophil mast cell progenitor, ProgB – B cell progenitor, RF – random forest, RFE – recursive feature elimination.

**Figure S8:**
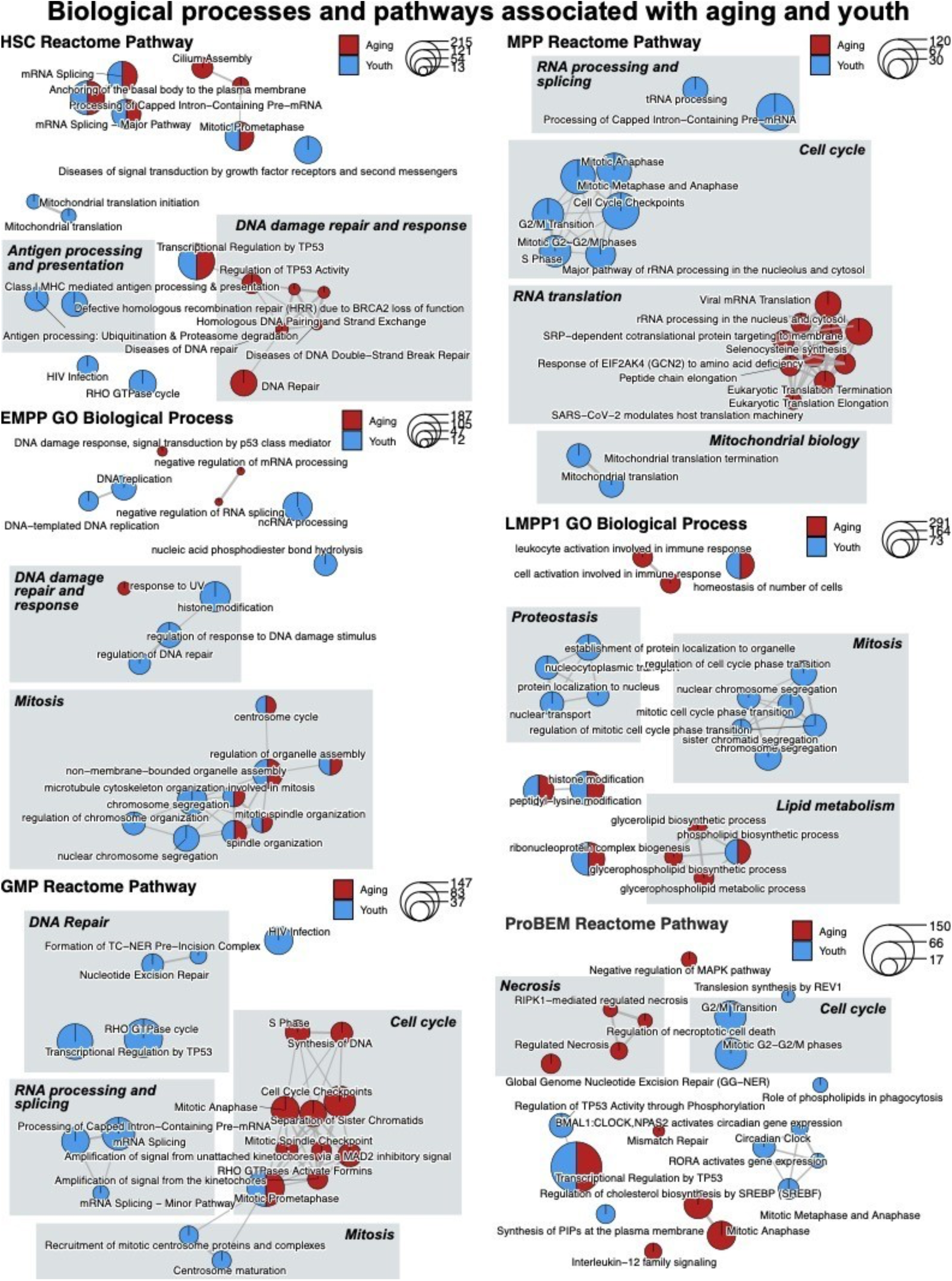
Celltype specific transcriptional changes are enriched for hallmark biological processes associated with aging. Select enrichment maps illustrating networks of overlapping gene sets enriched in genes associated with aging (red) and youth (blue). The gene sets analyzed were derived using a weighted spearman correlation analysis (see methods). Hallmark biological processes associated with aging are highlighted. EMPP – erythroid-biased MPP, GMP – granulocyte monocyte progenitor, HSC – hematopoietic stem cell, HSPC – hematopoietic stem and progenitor cell, LMPP – lymphoid-primed MPP, MPP – multipotent progenitor, ProBEM – basophil eosinophil mast cell progenitor.

Cells were sub-sampled from the integrated dataset (*n*=42,480) based on their annotations to perform a cell type specific analysis (*n*=2212-9553 cells per annotation). We also sub-sampled a similar number of cells (*n*=9274) across the entire dataset to resemble a comparable bulk sample (Bulk). The final cell type specific metrics are provided in Table S12. We first performed a weighted spearman correlation comparing gene expression within each cell type to both increasing age (aging) and decreasing age (youth) to identify important genes associated with aging and youth. These genes were then evaluated for important biological processes using an enrichment analysis (Figure 7C). A cell type specific approach improved the ability to identify significant terms compared to a similar analysis across all cell types (Bulk; Figure 7C-D). In aging HSCs, we observe enrichment in terms associated with antigen processing and presentation (Figure S8). In aging MPPs, we see enrichment in terms involved in TP53 signal transduction, DNA damage response, and ribosomal biology. In differentiated progenitors, including LMPP1s and GMPs, we observe enrichment in terms associated with inflammation, RNA processing and RNA splicing, DNA damage response and repair, and cell cycle. We were unable to identify terms associated with young age when analyzing all cell types together (Bulk; Figure 7C), however, by using a cell type specific approach we observe several cell type specific enrichment in terms associated with cell cycle, RNA biology, metabolism, and proteostasis (Figure S8).

Identifying important features that can predict individual cell age, can be an important tool for studying aging cells in single cell datasets. Although our ELN analysis identified important genes associated with aging, the gene sets comprise of >1614 genes (Table S13). We therefore asked if we could derive a simpler cell type specific aging signature using the integrated dataset to quantify cell age based on gene expression. To identify the most important cell type specific genes, we performed recursive feature elimination (RFE) using a random forest (RF) regression model and the caret R package on a training dataset comprising of 50% of the cells (Table S13). The cell type specific approach identified signatures containing 75-100 genes and improved our ability to predict cell age on the test dataset based on root mean square error (RMSE) and correlation (R^2^) (Figure 7D-E).

We then evaluated whether HSPCs in common blood cancers, including acute myeloid leukemia (AML) and pediatric acute lymphoblastic leukemia (ALL), express an aging phenotype. Published single cell datasets from patients with AML^31^ and pediatric ALL^67^ were labeled using our mixed model classifier. Cell types based on our annotations were sub-sampled and integrated with their healthy counterparts from the aging dataset (Figure 7B). We calculated cell type specific aging modules using the optimal features derived from our RFE analysis and compared them to healthy cells (Figure 7F). In both AML and pediatric ALL, which are aggressive myeloid and lymphoid blood cancers respectively, LMPP1s expressed higher levels of the LMPP1 aging score. There were too few ALL HSCs and GMPs for this analysis, however, HSC and GMP aging scores were not increased in AML. Prior work has shown that leukemia stem cell populations in AML are functionally similar to LMPPs^9^, and our results suggest that these cells may also express a more aged phenotype. Overall, these results strongly support a cell type specific analysis for identifying gene expression programs that are relevant in human aging and cancer.

## Discussion

In this study, we built a framework for characterizing the adult human MPP pool by generating combined scRNA-seq and scADT-seq data from normal human bone marrow donors and computationally identifying new HSPC cell states with distinct immunophenotypes (Figure 2), gene expression (Table S5), and chromatin accessibility (Figure 5). These cells were prospectively isolated using flow cytometry, and demonstrated unique functional properties based on complimentary *in vitro* and *in vivo* assays (Figure 3O). We show that within the classical MPP pool, only CLL1-CD69+ cells are capable of long-term engraftment and 7 lineage differentiation potential, whereas CLL1-CD69- and CLL1+ cells are erythroid-primed and myeloid-primed MPPs respectively with highly diminished long-term engraftment capacity. Within the classical LMPP sub-population, we show that CD2-CLL1-cells demonstrate high lymphoid potency, CD2+ cells express higher levels of cytotoxic genes and produce both lymphoid and myeloid cells, and CLL1+ cells contain primarily GMPs. Megakaryocyte production is restricted primarily to MPPs, erythrocyte production is enriched in EMPPs, lymphoid potency is greatest in LMPPs, and all evaluated HSPCs can produce granulocytes and monocytes (Figure 3O). It is worth noting that while the sub-populations purified by the new sorting strategy (Figure 2G, S3A-B) did enrich for the target cluster of interest, the final gates were not homogenous. Additionally, certain cells, ie MMPPs and LMPP1s, that were separated using differential surface marker expression had similar gene expression programs as measured by scRNA-seq, while having distinct functional output (Figure 3). This suggests that HSPCs exist in a gradient of cell states, and our sorting strategy primarily enriches for cells with certain characteristics rather than purifying for a homogenous population. This is an inherent limitation as the continuous nature of hematopoiesis is often not captured in discrete populations purified using FACS methods.

Regardless, the updated annotation and functional characterization of human HSPCs allowed us to build a valuable reference profile for evaluating external human and mouse data, which led to several key observations. Importantly, cell type specific approaches improved cell annotation transfer (Figures S4-6) which facilitated the subsequent analysis of external single cell datasets. We see that the cell states derived from our reference dataset are present in several external datasets and are analogous with external dataset annotations (Figure S6A). Of note, our ability to label cells in CB and mPB was highly diminished which again highlights the underlying molecular differences between different tissues in hematopoiesis (Figure S3C and S5C). Ultimately, our approach provides increased resolution for identifying HSPC states compared to published annotations (Figure S1E-F, S6A, and 7B).

Historically, human MPP populations have been less characterized compared to murine MPPs. This has led to difficulties in appropriately inferring the human relevance of observations made in mice. Our work provides new insights into the heterogeneity within the human MPP subpopulations, and our cross-species analysis with murine HSPCs identifies important relationships regarding homology between mouse and human HSPCs. We see that murine HSCs and more differentiated progenitors show a stronger one-to-one concordance with their analogous human states, compared to the MPPs and OPPs (Figure 6 and S7). These observations emphasize the underlying complexity and heterogeneity within both mouse and human MPPs, and the need for additional characterization of the murine MPPs.

Finally, we demonstrate the utility of our reference profile and mixed model classifier for studying human disease by performing a meta-analysis across 42,480 human HSPCs aged 0-75 (Figure 7). Using a cell type specific approach, we were able to nominate important biological processes in human aging and identified cell type specific associations for these derangements (Figure S8). Additionally, our approach allowed us to identify cell type specific aging scores that revealed interesting observations between the aging phenotype and blood cancers (Figure 7F) as AML and ALL LMPP1s, rather than HSCs, express a significantly higher aging score compared to healthy donors even in pediatric cases. We summarized the observations from this study into an updated model of adult hematopoiesis (Figure 4) which highlights the heterogeneity within the human MPP and OPP sub-populations and their associated functional capacities. Overall, our findings and methodologies provide new and important insights into adult MPP heterogeneity, and a valuable resource for future research in hematopoiesis.

## Limitations of the study

This study provides significant advancements in our understanding of adult human hematopoiesis, as we observe that cell states determined primarily by gene expression can be enriched within immunophenotypically defined sub-populations. It was therefore possible to use these surface markers to purify cells, although heterogeneous, to study their functional characteristics. However, our analysis was limited by the scarcity of primary adult human tissue and the diminished engraftment potential of adult BM compared to CB^21–24,55,68^. In fact, adult BM cells have not been reported to reliably produce multilineage engraftment over serial transplantations. We therefore had to evaluate populations of cells, rather than single cells, for our functional assays. This of course impairs our ability to definitively determine if our purified cells exist as a homogenous population or as a collection of cells with various functional capabilities even though this approach is consistent with historical advancements and refinements in HSPC immunophenotyping. Further, we were unable to show human cell engraftment in secondary transplants for either HSCs or CLL1-CD69+ MPPs (data not shown). Therefore, we did not relabel the MPPs as HSCs since positive multilineage engraftment over serial transplantations is required to define HSCs. Regardless of these limitations, we were able to observe significant differences amongst the purified cells which will be informative when designing and implementing translational research in human hematopoiesis. Future work will need to address the shortcomings inherent in our functional assays and continue to use complimentary cell sources, ie CB and mPB, and murine models. Although the reference profile provides an important framework for studying hematopoiesis, our computational analyses only nominate potential relevant biological processes. Future work will need to expand on these observations with the necessary mechanistic and functional correlates.

## Acknowledgements

We would like to thank Feifei Zhao for lab management and all members of the Majeti and Gentles labs for supporting this study. This work was supported by the National Institutes of Health (NIH) under awards 1R01HL142637 and 1R01CA251331 and the Stanford Ludwig Center for Cancer Stem Cell Research and Medicine (R.M.). A.J.G. was supported by the National Cancer Institute (NCI) of the NIH under awards R21CA238971, U01CA264611 and R01CA276828. A.E. was supported by the NCI under award F32CA250304, the Advanced Residency Training Program at Stanford, the American Society of Hematology (ASH) Scholar Award, and the Edward P. Evans MDS Young Investigator Award. A.C.F. was supported by a Stanford Graduate Fellowship, NSF Graduate Research Fellowship Program, and Stanford Lieberman Fellowship. T.K. was supported by the Leukemia and Lymphoma Society Special Fellow Award. S.H. was supported by the ASH Medical Student Physician Scientist Award. B.A.L. was supported by the NIH under award K99CA276901. Y.K. was supported by Stanford Immunology Baker Fellowship and by KFAS Overseas PhD Scholarship from Korea Foundation for Advanced Studies. M.H.L. was supported by the Blavatnik Family Fellowship. A.A. was supported by ASH HONORS Award. Some of the computing for this project was performed on the Sherlock cluster. We would like to thank Stanford University and the Stanford Research Computing Center for providing computational resources and support that contributed to these research results. The content is solely the responsibility of the authors and does not necessarily represent the official views of the National Institutes of Health.

## Author contributions

Conceptualization, A.E. and R.M.; Methodology, A.E., A.J.G., A.M.N and R.M.; Investigation, A.E., Y.N., A.C.F., X.H., S.R., M.N., A.A.; Writing – Original Draft, A.E. and R.M.; Writing – Review & Editing, A.E., A.J.G., A.M.N., R.M.; Funding Acquisition, R.M.; Supervision, A.C.F., T.K., B.A.L., Y.K., D.K., M.H.L., A.J.G., A.M.N., and R.M.

## Declaration of interests

R.M. is on the Advisory Boards of Kodikaz Therapeutic Solutions, Orbital Therapeutics, Pheast Therapeutics, and 858 Therapeutics. R.M. is a co-founder and equity holder of Pheast Therapeutics, MyeloGene, and Orbital Therapeutics. S.R. and M.N. previously worked at BD Biosciences. All other authors declare no competing interests.

**Graphical Abstract: Multi-modal framework for isolating and studying HSPCs in adult hematopoiesis.** Schematic illustrating the methods used to characterize adult HSPCs. Bone marrow cells from three healthy donors underwent single cell whole transcriptome analysis (scRNA-seq) and surface marker quantification (scADT-seq). Data were first clustered and annotated based on marker genes. New immunophenotypes were derived computationally to purify each HSPC cluster and converted to a sorting strategy using flow cytometry. HSPCs were subsequently purified from both adult and cord blood donors and studied using both *in vitro* and *in vivo* assays. Based on their unique molecular and functional properties, a new reference profile of adult HSPCs was used to perform an integrative analysis on chromatin accessibility, cross species homology, and celltype specific changes with aging. EMPP – erythroid-biased MPP, GMP – granulocyte monocyte progenitor, HSC – hematopoietic stem cell, HSPC – hematopoietic stem and progenitor cell, LMPP – lymphoid-primed MPP, MPP – multipotent progenitor, MPPP – myeloid-biased MPP, NMF – non-negative matrix factorization, ProBEM – basophil eosinophil mast cell progenitor, ProgB – B cell progenitor.

## STAR*METHODS

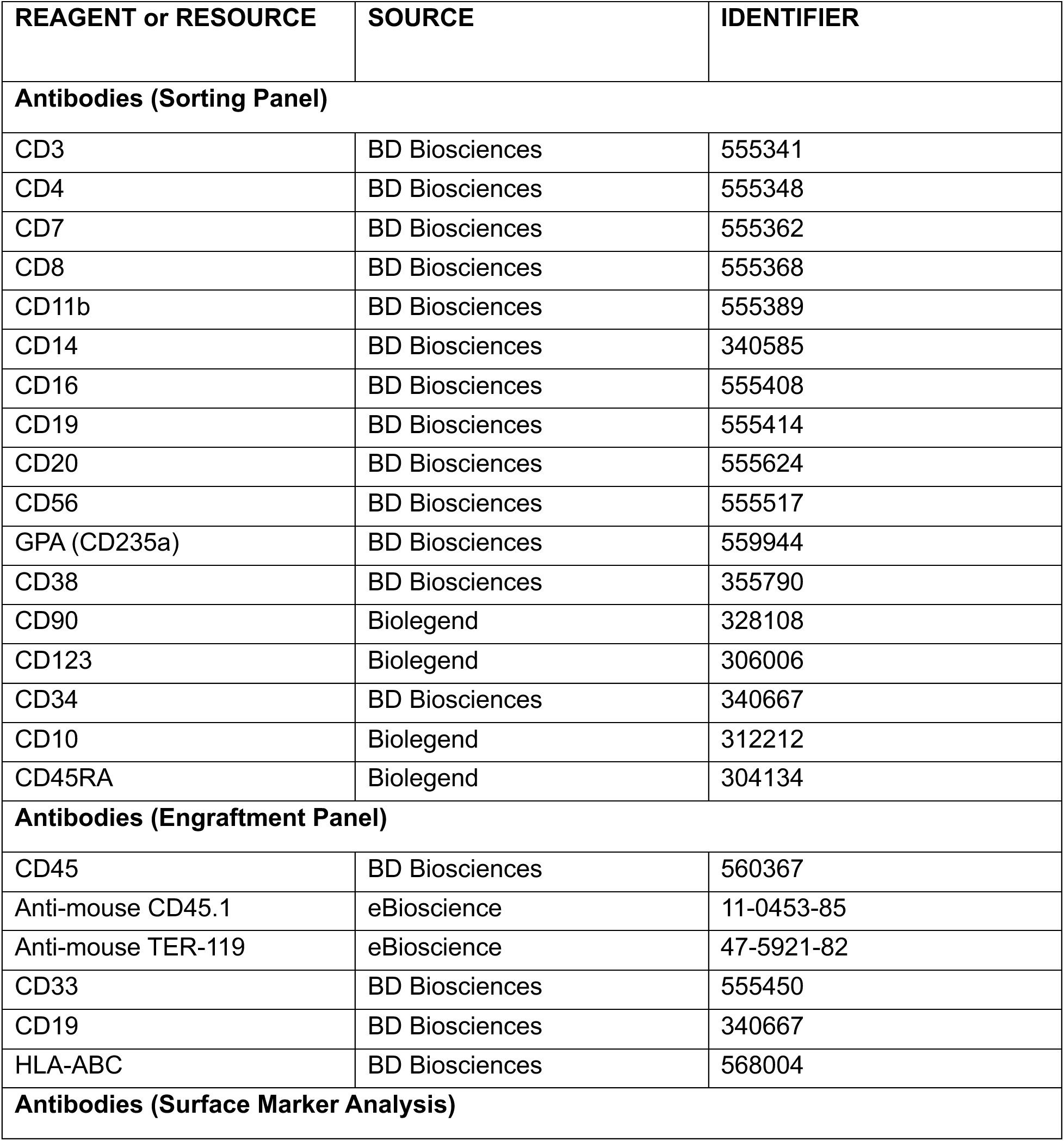
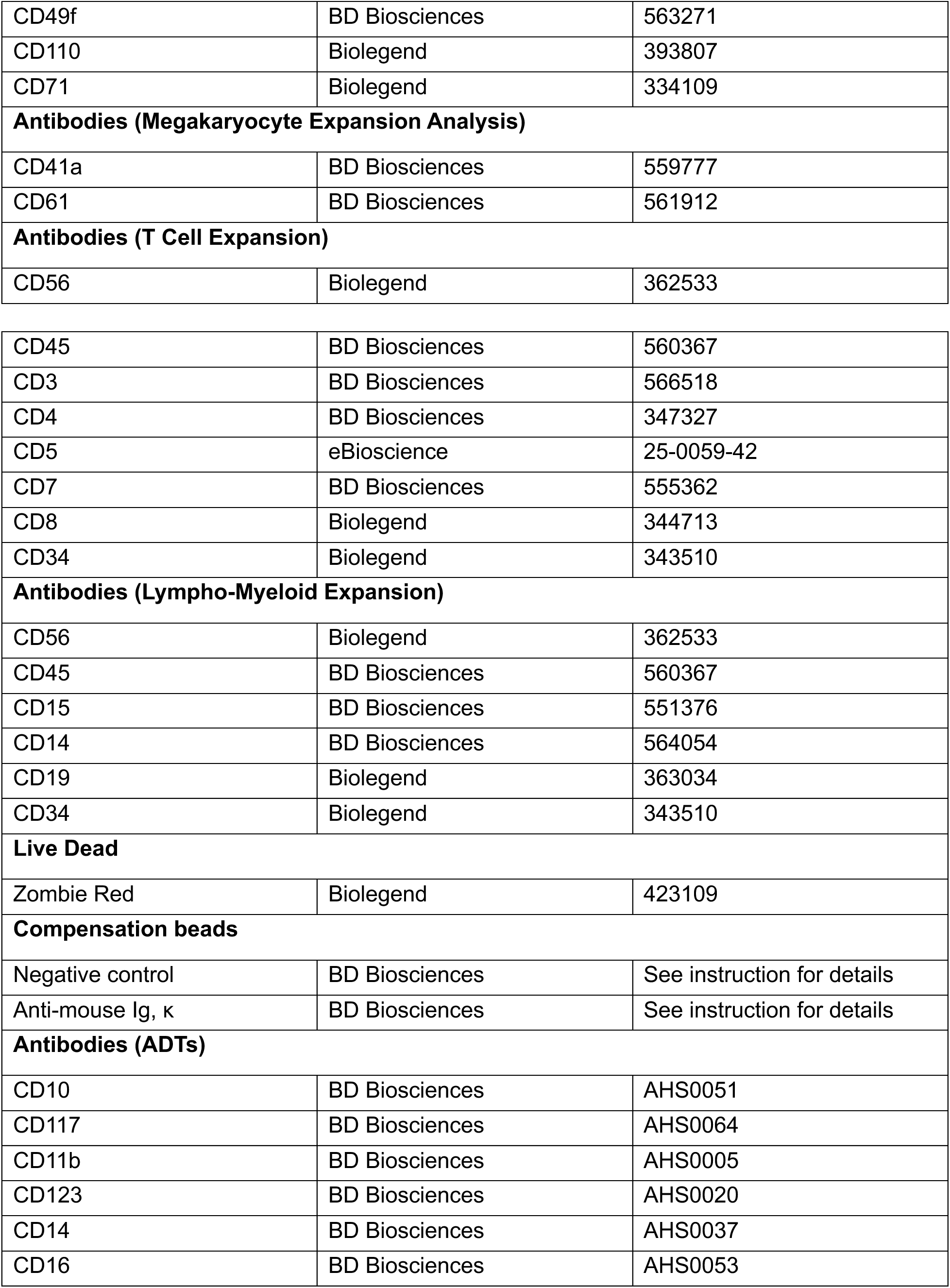
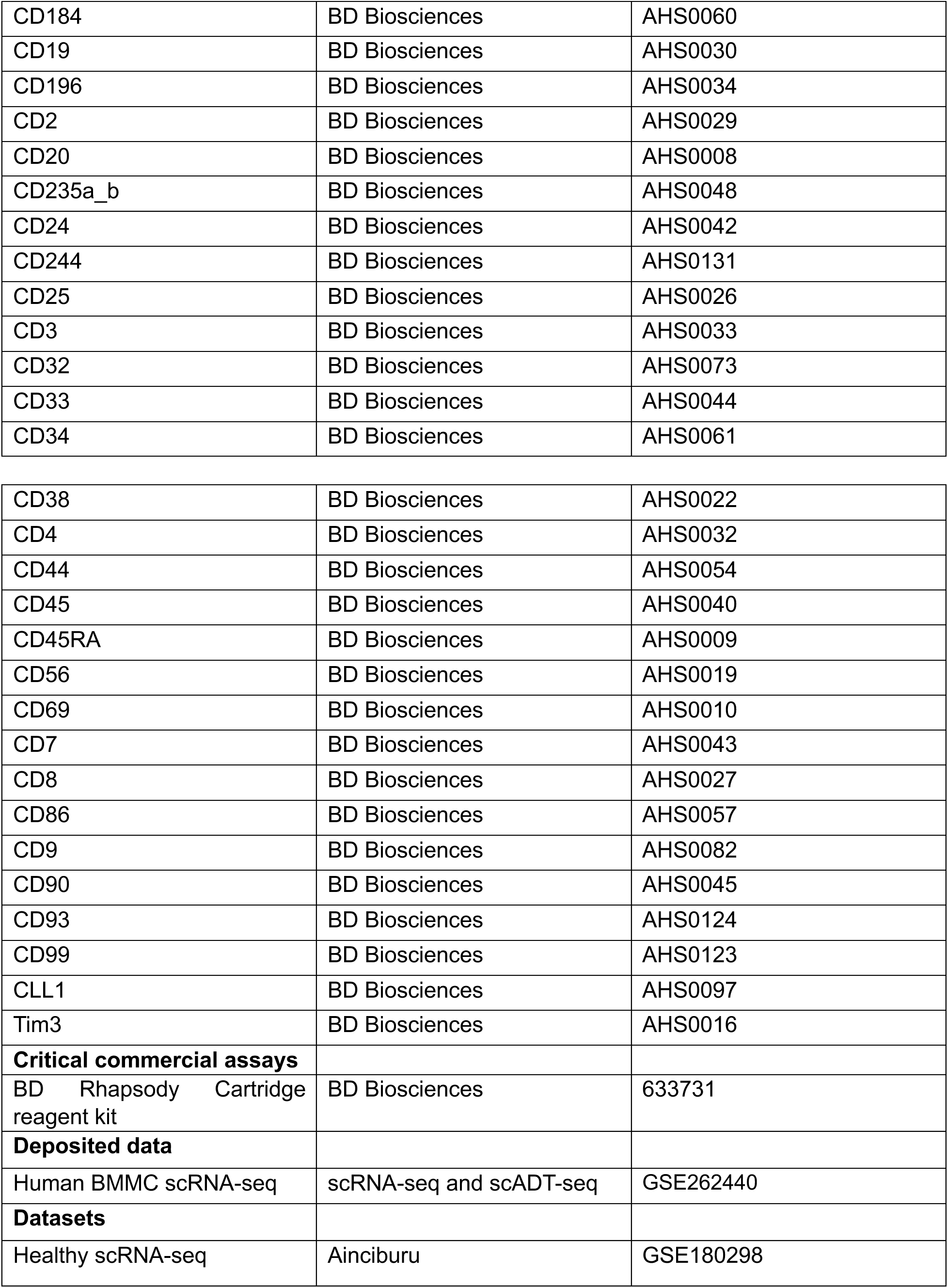
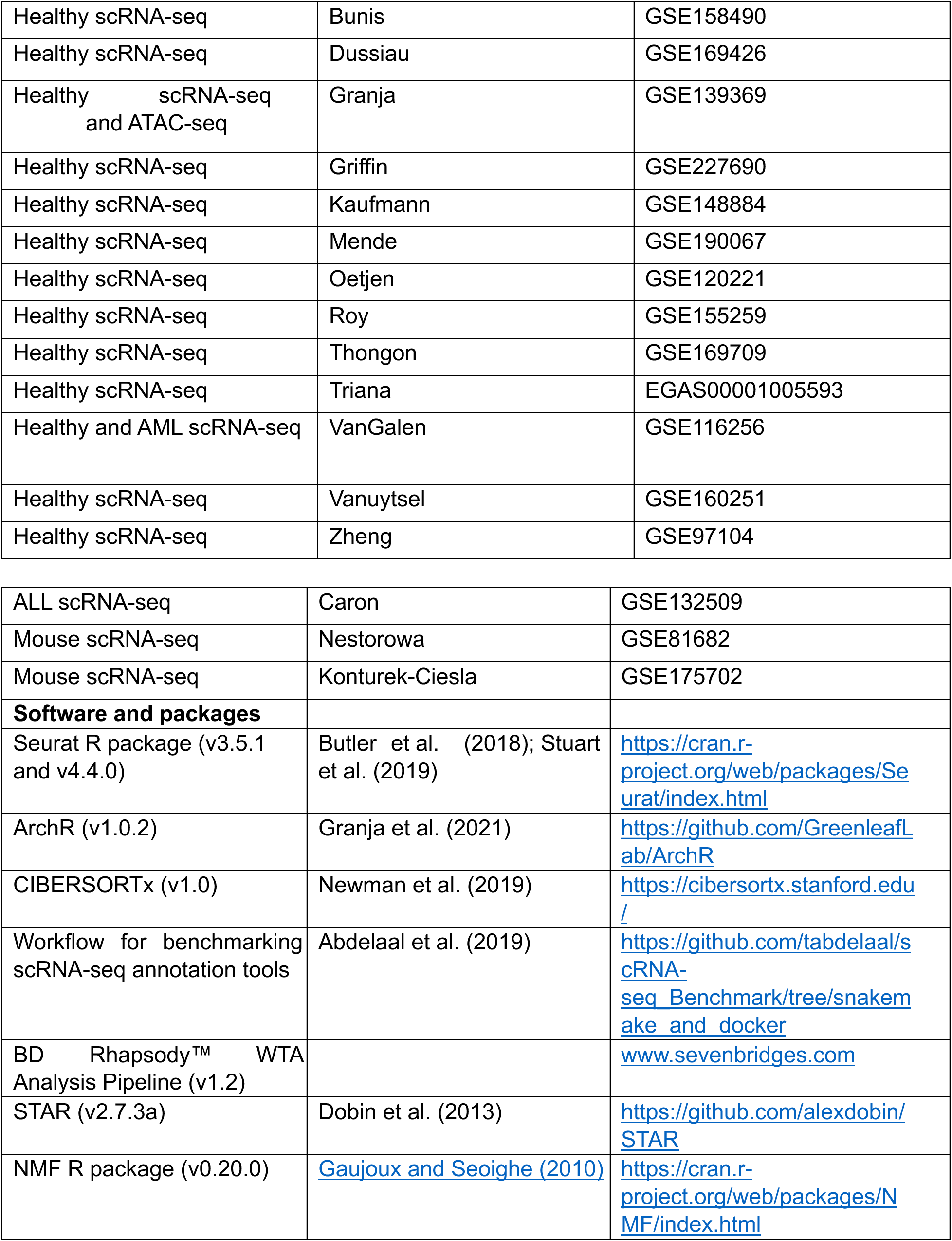
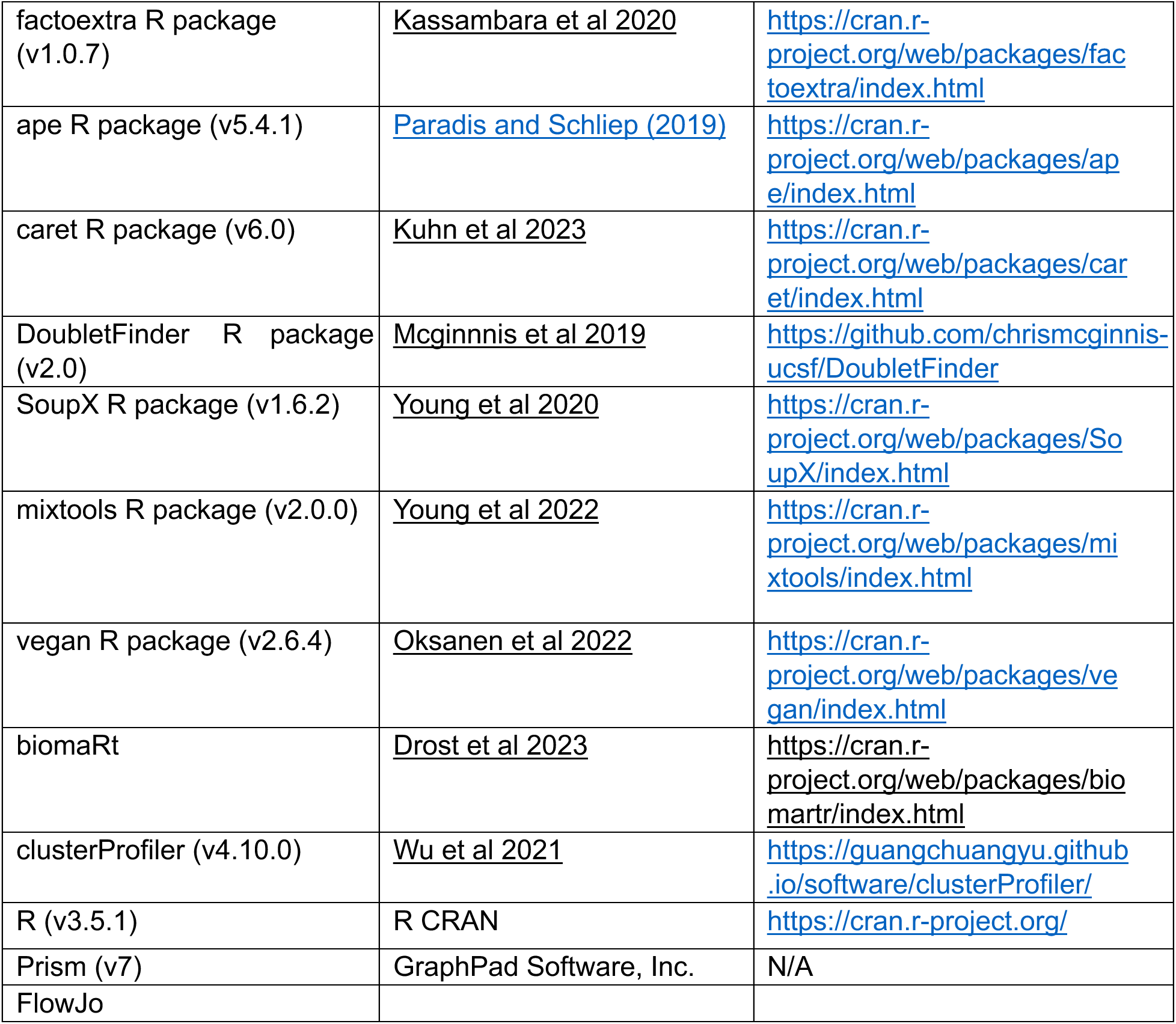
KEY RESOURCE TABLE

## RESOURCE AVAILABILITY

### Lead contact

Further information and requests for resources should be directed to and will be fulfilled by the lead contact, Ravindra Majeti.

### Materials availability

This study did not generate new unique reagents.

### Data and Code Availability

The data generated in this paper was deposited in GEO under accession GSE262440.

## METHOD DETAILS

### Description of healthy adult donors

Bone marrow mononuclear cells were obtained from healthy donors with informed consent and compliance with relevant ethical regulations (AllCells and STEMCELL Technologies). All healthy cells used in this study were cryopreserved, and subsequently thawed by the dropwise addition of IMDM (GIBCO, cat. no. 12440-061) containing 20% fetal bovine serum (FBS; Sigma, cat. no. F1051). Characteristics of each donor, including age and sex, are available in Table S1.

### Single cell captures for whole transcriptome and surface marker quantification

Cryopreserved adult bone marrow cells were thawed and stained with annexin V (Thermo Fisher Scientific) per protocol for 15 min. Cells were resuspended with DAPI and annexin V/DAPI negative cells were sorted for subsequent ADT staining and single cell capture per established protocols (BD Rhapsody™). The BD Rhapsody™ workflow is a nanowell based single cell capture system with established protocols for generating single cell whole transcription and ADT cDNA libraries. Briefly, an ADT cocktail (1uL per ADT) was prepared in a total of 100 uL of buffer. Cells were first washed and then labeled with the ADT cocktail for 30 min at 4C, washed 3 times, resuspended in sample buffer (BD Rhapsody™ Cartridge reagent kit), and captured with the BD Rhapsody™ multiomic single cell system using the manufacturer’s instructions^69^. ADT and whole transcriptome libraries were generated per manufacture’s protocol, evaluated by qubit and bioanalyzer, pooled and sequenced using NextSeq500 or Illumina Novaseq S2 (Illumina). Sequencing depth was determined based on manufacturer’s recommendations.

### Sample processing, integration, and quality control

Single cell datasets are inherently noisy, as multiplets and pre-apoptotic cells can significantly confound downstream analysis^70,71^. Further, non-specific ADT binding mediated by not only antibody cross-reactivity and poor input cell quality but also complex biological interactions^70–72^ can further complicate analysis of multimodal single cell datasets. Therefore, we applied stringent quality control that incorporates the basic Seurat workflow for multimodal data processing and integration^73,74^, ambient RNA and ADT removal^75,76^, doublet discrimination using DoubletFinder^77^, and elimination of cells expressing mutually exclusive lineage markers (Figure 1A, S1A-B).

Fastq files generated from the BD Rhapsody™ workflow were processed on Seven Bridges (https://www.sevenbridges.com) using the standard analysis pipeline (BD Rhapsody™ WTA Analysis Pipeline). The unique molecular identifier (UMI) count matrices were imported into R (v3.6.2) and processed using Seurat (v3.5.1 and v4.4.0)^74^. Each domain, ie mRNA and ADT counts, was normalized using log-normalization or centered using log ratio normalization (CLR) respectively. A third concatenated dataset was created by merging normalized counts for specific down-stream analyses. Poor quality cells were removed based on total RNA content and percent mitochondrial content using the top 1% expression level. Doublet discrimination was first performed using DoubletFinder with standard metrics^77^. Doublets were removed and gene and surface marker expression were corrected for ambient noise using SoupX^75^. DoubletFinder was used to filter doublets rather than a maximum RNA content filter due to the inherent variability of RNA content and celltype^78^. Mutually exclusive lineage markers (Table S4) were subsequently used to remove potential heterotypic doublets, and poor-quality cells resulting from non-specific ADT binding^71^. These strict filtering thresholds removed approximately 20% of cells rather than the expected 8-9% based on loading metrics (Table S2). Of note, removing cells based on doublet discrimination would have removed only 7% of cells. We also see that cells removed based on conflicting lineage marker expression had significantly higher ambient ADT signal (rho) as measured by SoupX, despite having similar mitochondrial and RNA content (Figure S1B). The filtered cells were subsequently integrated by sample using canonical correlation analysis (CCA) and the resulting integrated matrix containing the top 5000 mRNA and ADT features was used for downstream computational analyses (Figure S1A).

### Clustering and diversity analysis

Cells were computationally purified using standard and new HSPC gating strategies (Table S4) with the CellSelector function in Seurat. If needed, gating thresholds were determined by modeling the expression of each ADT using mixture models (*k*=2-3) and the expectation maximation algorithm with mixtools^79^. The gap statistic was calculated using the integrated matrix described above within designated sub-population using the factoextra R package (nstart=25, 100 iterations, d.power=2, Tibs2001SEmax, and K-means clustering)^37^. Sub-population purity was assessed using the shannon diversity index and the vegan R package^80^.

### NMF and Louvain clustering

Louvain clustering was performed using Seurat and the resolution was adjusted to achieve target cluster number. Clustering using the non-negative matrix factorization (NMF) approach was performed as previously described^39^ using the NMF R package^40^. Briefly, the top 2000 variable features (31 ADTs; Table S4) were identified using the normalized count matrix from the integrated assay containing both mRNA and ADT expression. After nonnegative transformation, the NMF algorithm was performed for k=2-20 clusters. Optimal cluster number was estimated using the cophenetic index. Initially, the data was over-clustered and annotated using marker features, and clusters with low numbers (n<10) were removed. We used CIBERSORTx^41^ to identify similar clusters by first splitting the data into a test and training set (1:2 split), and a signature matrix was created using the training dataset. Artificial bulk transcriptomes were created using the test dataset by subsetting cells based on their cluster label, converting counts to transcripts per million (TPM), and summing counts across all cells. We deconvolved these artificial bulk transcriptomes using the signature matrix, and poor performing clusters were collapsed with the closest mislabeled cluster. The resulting set of clusters were evaluated for established lineage or cell defining marker expression and sub-annotated accordingly (higher resolution immunophenotype; Table S4). With Seurat, differentially expressed genes and surface markers were determined using the FindAllMarkers and the MAST test, and cell cycle scores were calculated using the CellCycleScoring function.

### New surface marker identification

We performed an iterative differential surface marker analysis to identify a specific immunophenotype for each newly identified HSPC sub-population. HSCs, MPP1-3s, LMPP1-2s, GMPs, and ProBEMs were isolated from the original dataset. Differential ADTs were first determined across all cell populations. Differential markers were evaluated manually, and cells were computationally purified based on informative markers and then re-evaluated for differential markers (Table S4). Candidate features were subsequently evaluated using FACS to identify markers with sufficient dynamic range to efficiently purify each sub-population (data not shown).

### Animal care

All mouse experiments were conducted in accordance with a protocol approved by the Institutional Animal Care and Use Committee (Stanford Administrative Panel on Laboratory Animal Care #22264) and in adherence with the U.S. National Institutes of Health’s Guide for the Cand Use of Laboratory Animals.

### Fluorescence-activated cell sorting

Thawed cells were washed with FACS buffer (PBS, 2% FBS, 2 mM EDTA) and stained with antibodies for 30 minutes on ice in 50 uL total volume. Cells were then stained with Zombie Red for viability assessment for 5 minutes and subsequently washed prior to analysis. All cell sorting steps were confirmed using post-sort purity analyses. The antibodies used for respective assays are provided in Table S4. Flow cytometry was performed on a FACSAria II (Becton Dickinson). Gates were drawn by internal positive and negative controls using healthy cells and validated by back-gating on marker positive events. Positive gates for CD49f expression were based on peak expression for the HSCs from a particular donor, and the same gate was used for other samples from the same donor. This allowed us to quantify relative CD49f expression compared an internal control. Gates drawn for all other markers were consistent across all samples (Figure S2).

### Single cell colony formation assay

Single HSPCs were sorted into 96 well plates containing 120 uL of Methocult H4034 methylcellulose media (STEMCELL Technologies). Cells were allowed to differentiate for 12 days and colonies were morphologically assessed. The ratio of colonies per total cells seeded is presented for each celltype.

### Megakaryocyte expansion assay

Purified HSPCs were plated in 96 well plates containing megakaryocyte differentiation medium (StemSpan SFEM II with human low-density lipoproteins (STEMCELL Technologies) and Megakaryocyte Expansion Supplement (STEMCELL Technologies)). Megakaryocyte production was assessed at 7 days by staining cells for CD41a and CD61 expression. Megakaryocyte number was normalized to live cells and 1000 cells seeded.

### Xenotransplantation assays

Purified HSPCs were transplanted into the right femur of sub-lethally irradiated NSG mice (6-8 weeks, male and female, 200 rad 2-24 hours pre-transplant)^44^. Mice were harvested at 17-18 weeks by crushing the pelvis and bilateral femurs and tibias. Cells were filtered and stained using the engraftment panel (Table S4) to assess both engraftment and lineage. Human to mouse chimerism was calculated based on hCD45+HLA-ABC+ cells relative to mouse CD45+mTer119-cells. Chimerism >0.1% was considered positive for long term engraftment.

### OP9 and OP9-hDL4 maintenance

OP9 and OP9-hDL4 cells were gifted from the Bendall Lab at Stanford University. Cells were cultured in fresh OP9 media (MEMα (Gibco12561056), 15% FBS, 1% L-glutamine, and 1% Penicillin-Streptomycin (Gibco)). Cells were split 1:4 or 1:5 after reaching 90% confluency.

### OP9 co-culture lymphoid-myeloid cell differentiation assay

OP9 cells were seeded in a 96 well plate (2500 cells/well) in 37C, 5% CO2 incubation. Twentyfour hours after plating, purified HSPCs were added to each well and expanded with 20 ng/ml SCF, 10 ng/ml FLT3L, 10 ng/ml G-CSF, 10 ng/mL IL-2 (all from Peprotech, London, UK), 10 ng/mL IL-15 (STEMCELL Technologies), and 0.2 uM DUP697 (Cayman Chemicals) in OP9 media (SGF15 media). Cells were dissociated and transferred to new plates with fresh OP9 cells and SGF15 media weekly. Cells were analyzed by flow cytometry at weeks 1, 2 and 3. Lineage output was assessed by staining cells for CD45, CD34, CD15, CD14, CD19, and CD56 according to immunophenotypes described in Table S4. Cell output was normalized to hCD45 content for lineage output quantification and both hCD45 content and cells seeded for production quantification.

### OP9-hDL4 co-culture T cell differentiation assay

OP9-hDL4 cells were seeded in a 96 well plate (2500 cells/well) in 37C, 5% CO2 incubation. Twenty-four hours after plating, purified HSPCs were added to each well and expanded with 10 ng/ml SCF, 5 ng/ml FLT3L and 5 ng/ml IL-7 (Peprotech, London, UK) in OP9 media (SF7 media). Cells were dissociated and transferred to new plates with fresh OP9-hDL4 cells and SF7 media weekly. Cells were analyzed by flow cytometry at weeks 3 and 4. Lineage output was assessed by staining cells for CD45, CD34, CD7, CD5, CD3, CD4, and CD8 according to immunophenotypes described in Table S4. Cell output was normalized to hCD45 content for lineage output quantification and both hCD45 content and cells seeded for production quantification.

### Integrated single cell ATAC sequencing analysis

The scRNA-seq data was integrated with published scATAC-seq data from adult donors with ArchR using the developers’ recommendations unless otherwise stated^30,50^. Briefly, arrow files were generated using published read fragment data, low quality cells and doublets were removed, a tile matrix was created, and dimensional reduction was performed using latent semantic indexing. Gene activity scores were calculated using local accessibility of primarily the promoter and gene body. The addGeneIntegrationMatrix function was used to perform a constrained integration (Table S11). Pseudobulk replicates were constructed using the addGroupCoverages function, peak calling was performed using the addReproduciblePeakSet, and a peak matrix was created using addPeakMatrix. Differentially opened peaks were determined using addMarkerFeatures, motif annotations were added using the addMotifAnnotations function and the Vierstra database, deviations were calculated using addDeviationMatrix and getGroupSE, and footprints were identified using getFootprints. Correlated gene expression using the gene integration matrix and motif deviations were calculated using the correlateMatrices function to identify putative regulators. Trajectories were added using the addTrajectory function and motif deviations and gene expression were correlated across pseudotime to identify differential regulators (Table S9).

### Deriving a mixed model classifier for cell label transfer

Counts from each respective dataset (Table S7) were processed and integrated as described above. Code provided by Abdelaal et al^52^ was adapted to evaluate classifiers based on several models: CHETAH^81^, scID^82^, RF^83^, singleR^84^, singleCellNet^85^, scmapcell^86^, SVM (linear kernel with and without rejection)^87^, and scPred (MDA, NNet, avNNet, GLM, SVM with radial kernel)^88^. The intra-dataset analysis was performed as described by the authors using 10-fold CV, and default parameters for each classifier. The top models were identified based on F1 scores, and proportion of cells labeled and evaluated using a celltype specific approach with scPred with default parameters. Top performing models by celltype was used for the final mixed-model classifier. For the inter-dataset analysis, we first held out 10% of the data, and the remaining 90% of data was used to train the model. We trained our mixed model classifier using the training dataset which was used to assign labels on the external dataset, ie the query dataset. We then used the labeled query dataset to retrain the mixed-model classifier which then assigned labels onto the left-out test dataset from our reference. This approach allowed us to indirectly evaluate the performance of the model since the test dataset contains the true cell labels. This modified 10-fold CV approach was used to evaluate classifier performance. For final cell label assignments, we used the entire reference dataset and a probability threshold of 0.75 rather than the default of 0.5 to label the query dataset, and unassigned cells were removed from the subsequent analysis.

### Cross species analysis

Gene aliases were relabeled using gene ortholog-to-ortholog conversions in R with biomaRt^89^ and murine single cell count matrices were subset to contain only conserved orthologs^74^. Non-HSPCs were removed in the mouse data to ensure that comparable cell states were being evaluated in the downstream analysis. Count tables were processed in Seurat as described above. Murine cells were labeled using the mixed model classifier, and the data were clustered using the Louvain algorithm. Conserved features were identified using the FindConservedMarkers function in Seurat, and murine and human cells were integrated using only conserved markers with Seurat CCA. Co-occurrence heatmaps were generated for each integrated data set, and average gene expression per cluster was determined using the AverageExpression function in Seurat to perform a gene expression correlation analysis.

### Aging analysis

Datasets with both age and celltype annotations were included in this analysis (Table S12) and processed as described above. Samples were categorized based on age (Table S12). Each dataset was labeled using the mixed model classifier using the reference profile annotation described in Figure 1D. Cells were subset by celltype and integrated as described above. To identify important genes associated with aging, gene expression was correlated with age using a spearman correlation weighted by cells per age (Table S9). Important genes were selected based on significance (adjusted p-value <0.05 based on the Benjaminid-Hochberg procedure) < 0.05. GO term over representation analysis using these important genes was performed using clusterProfiler in R^90^. To derive celltype specific aging signatures, data were split into test and training sets (1:1 split) and RFE was performed using caret with RF regression models on the training dataset. Informative features based on RFE was used to predict age in the test dataset and root mean squared error (RMSE) and correlation (R2) was calculated. To study aging signatures in human blood cancers, external datasets (Table S12) were processed as described above and module scores based on informative features from the RFE analysis (Table S13) using the AddModuleScore function in Seurat.

### Quantification and Statistical Analysis

Details of statistical tests performed can be found in figure legends or materials and methods. For significance testing, a t-test (2-tailed distribution and two-sample equal variance) was used for comparing 2 conditions or a one-way ANOVA with Tukey’s multiple comparison adjustment when comparing >2 conditions unless otherwise stated. Results with adjusted p-values <0.05 were considered significant. All statistical analyses were performed in R or Graphpad Prism.

